# Can single-neuron frequency tuning in human auditory cortex be quantified through fMRI adaptation?

**DOI:** 10.1101/2022.01.06.475208

**Authors:** Julien Besle, Rosa-María Sánchez-Panchuelo, Susan Francis, Katrin Krumbholz

## Abstract

Measuring neuronal frequency selectivity in human auditory cortex may be crucial for understanding common auditory deficits such as speech-in-noise difficulty. Non-invasive methods measure aggregate responses of large populations of neurons and therefore overestimate single-neuron tuning selectivity. Here we explore whether cortical frequency selectivity can be estimated through fMRI adaptation. Using ultra-high-field (7T) BOLD-fMRI and individualized functional parcellation of auditory cortex, we measured the suppression (or adaptation) of primary and non-primary cortical responses to a high-frequency (3.8 kHz) probe sound as a function of the frequency of a preceding adaptor sound (ranging from 0.5 to 3.8 kHz). The degree of frequency tuning of the adaptation effect strongly depended on the temporal structure of the adaptor. Suppression by a single 200-ms adaptor showed little or no tuning, despite clear frequency tuning of the responses to the different adaptors. In contrast, suppression by multiple (four) 50-ms adaptor bursts was clearly tuned, with greater frequency selectivity than the adaptor response tuning, suggesting that fMRI adaption to multiple adaptors may reflect the frequency tuning of the underlying neuronal response. Importantly, adaptation tuning differed between primary and non-primary regions, suggesting a local suppression effect, rather than inheritance from upstream subcortical structures. Using a computational model of fMRI adaptation in a tonotopically-organized neuronal array, we identify key factors determining the relationship between observed fMRI adaptation tuning and the frequency selectivity of the underlying neuronal response. Using this model, we derive a plausible range for the frequency selectivity of individual neurons in each region of auditory cortex.

## Introduction

Frequency selectivity is a fundamental organizing principle of the auditory system, and accurate frequency representation is crucial for auditory perception. Animal work has suggested that neuronal frequency tuning changes from the cochlea to the cortex, but there is no consensus yet as to whether cortical frequency tuning is broader or, in fact, finer than cochlear frequency tuning [1, 2]. In humans, cochlear frequency tuning can be estimated either behaviourally, using notched-noise masking [3-5], or from otoacoustic emission latencies [6], but there is currently no accepted measure of neuronal frequency tuning in the human auditory cortex. Here, we explore whether cortical frequency tuning can be estimated through fMRI adaptation.

Previous fMRI studies have estimated population frequency tuning properties in human auditory cortex by measuring voxelwise blood oxygen level-dependent (BOLD) responses to different frequencies [7-10]. Voxelwise response tuning estimates, whilst useful for locating primary (or “core”) auditory areas [thought to exhibit greater frequency selectivity than surrounding secondary or “belt” areas: 7, 10], reflect the aggregate tuning properties of tens to hundreds of thousands of neurons [11, 12], and thus overestimate the frequency selectivity of individual neurons.

The two main methods that have been proposed for inferring single-neuron tuning properties from fMRI responses are multivariate pattern analysis (MVPA) and fMRI adaptation. MVPA uncovers subtle differences in the spatial distribution of voxelwise responses to different stimuli [13]. Although these differences are thought to reflect differences in within-voxel distributions of neuronal selectivity, MVPA does not provide a direct estimate of neuronal tuning curves. In contrast, fMRI adaptation could potentially be used to infer the shape and width of neuronal tuning curves. fMRI adaptation measures the degree of suppression, or adaptation, of the response to a probe stimulus caused by a preceding adaptor stimulus. Adaptation is maximal when adaptor and probe are identical, but if the adaptor and probe differ on a given stimulus feature, such as frequency in hearing or stimulus orientation in vision, adaptation may be reduced, and the amount of reduction in adaptation is thought to reflect the degree of neuronal tuning to the relevant feature [14, 15]. Adaptation paradigms have been extensively used in visual fMRI studies to demonstrate neuronal selectivity to features such as orientation or motion direction within different visual cortical areas [e.g., 16, 17, 18], but have only rarely been used to obtain quantitative estimates of neuronal tuning width [19-21]. This is because the relationship between fMRI adaptation tuning and neuronal tuning is complex and still poorly understood [22, 23]. Whilst often assumed to reflect neuronal tuning properties, fMRI adaptation likely also comprises hemodynamic components, the tuning properties of which likely differ from the neuronal tuning properties [24]. Even at the neuronal level, adaptation tuning may differ from response tuning, and has been reported to be either more widely tuned [e.g. 25], tuned to different stimuli [e.g. 26], or even more narrowly tuned if adaptation is inherited from more sharply tuned downstream structures [23, 27].

The current study aimed to explore and disentangle these factors in the case of frequency tuning in human auditory cortex. Using a simple adaptor-probe paradigm, we measured both response and adaptation tuning for frequency, in both primary and non-primary auditory fields, delineated through individualized in-vivo parcellation based on tonotopic gradient reversals [7]. As previous results have suggested differences in adaptation tuning properties between transient response components at stimulus onset and subsequent sustained components [25], we presented adaptors as either single-onset or multiple-onset bursts (with equated total energy). Using a computational model of tonotopically-organized auditory cortex, we relate the measured BOLD-response and fMRI-adaptation tuning properties to possible underlying neuronal and hemodynamic factors and derive a range of plausible neuronal tuning width estimates for both primary and non-primary auditory regions. @@@

## Results

### FMRI adaptation in the human core and belt auditory cortex is frequency-specific

To investigate the frequency-specificity of fMRI adaptation, we measured BOLD responses to adaptor and probe stimuli presented either in isolation (A and P responses), or as adaptor-probe pairs (AP response) (Fig. 1). The adaptor frequency was varied between 0.25 and 6.01 kHz for adaptors only (A), and between 0.51 and 3.84 kHz for adaptor-probe pairs (AP), whilst the probe frequency was fixed at 3.84 kHz (Fig. 1A). In addition, we manipulated the temporal properties of the adaptor stimulus: for the first five participants, the adaptor was a single 200-ms stimulus (single-onset adaptor condition, Fig. 1B), while, for the next 6 participants, it was a train of four identical 50-ms stimuli, separated by 30-ms silent gaps (multiple-onset adaptor condition, Fig. 1C). In either case, the probe stimulus was a 50-ms stimulus.

**Figure 1:**
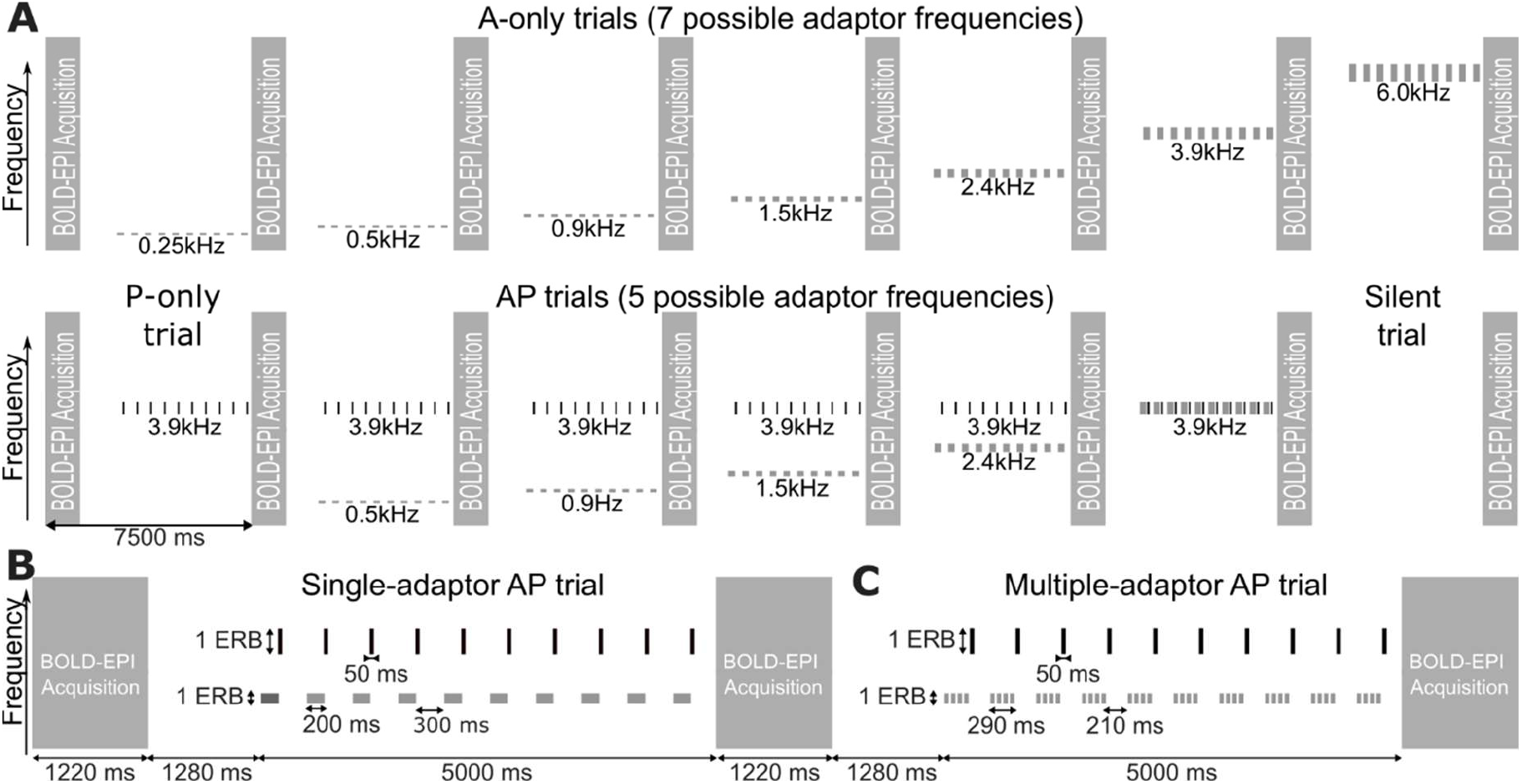
Stimulus presentation and fMRI acquisition timing. A. Timelime of 15 example sparse fMRI trials showing all seven adaptor (A) trials, all five adaptor + probe (AP) trials, one probe (P) trial and one null (silent) trial. The long vertical bars represent the fMRI volume acquisitions. The short bars show the stimulus trains presented in between acquisitions. All stimuli were narrowband noises whose center frequency and tuning width are indicated on the vertical axis (representing a linear frequency scale). B&C. Detailed timeline of a single AP trial containing either single-onset 200-ms adaptors (B; first 5 participants) or multiple-onset 50-ms adaptors (C; last 6 participants). A and P trials followed the same timeline, with either the probes or the adaptors removed from the train.

We first used the A responses at the seven frequencies (averaged across single-onset and multiple-onset adaptors) to derive tonotopic maps in each hemisphere of each participant (Fig. 2A), which, in turn, were used to define four regions of interest (ROIs) corresponding to the core (primary) and belt (non-primary) auditory cortex. In each individual hemisphere, we identified two mirror-reversed tonotopic gradients overlapping the core region [as marked by increased intracortical myelin and frequency selectivity in and around Heschl’s gyrus, Fig. 2B,C; see ref. 7]: the anterior gradient ROI (overlapping most with the core) and the posterior gradient ROI (Fig. 2D). The other two ROIs, henceforth referred to as anterior and posterior belt ROIs comprised all auditory-responsive voxels located either anteriorly or posteriorly to the two gradient ROIs.

**Figure 2:**
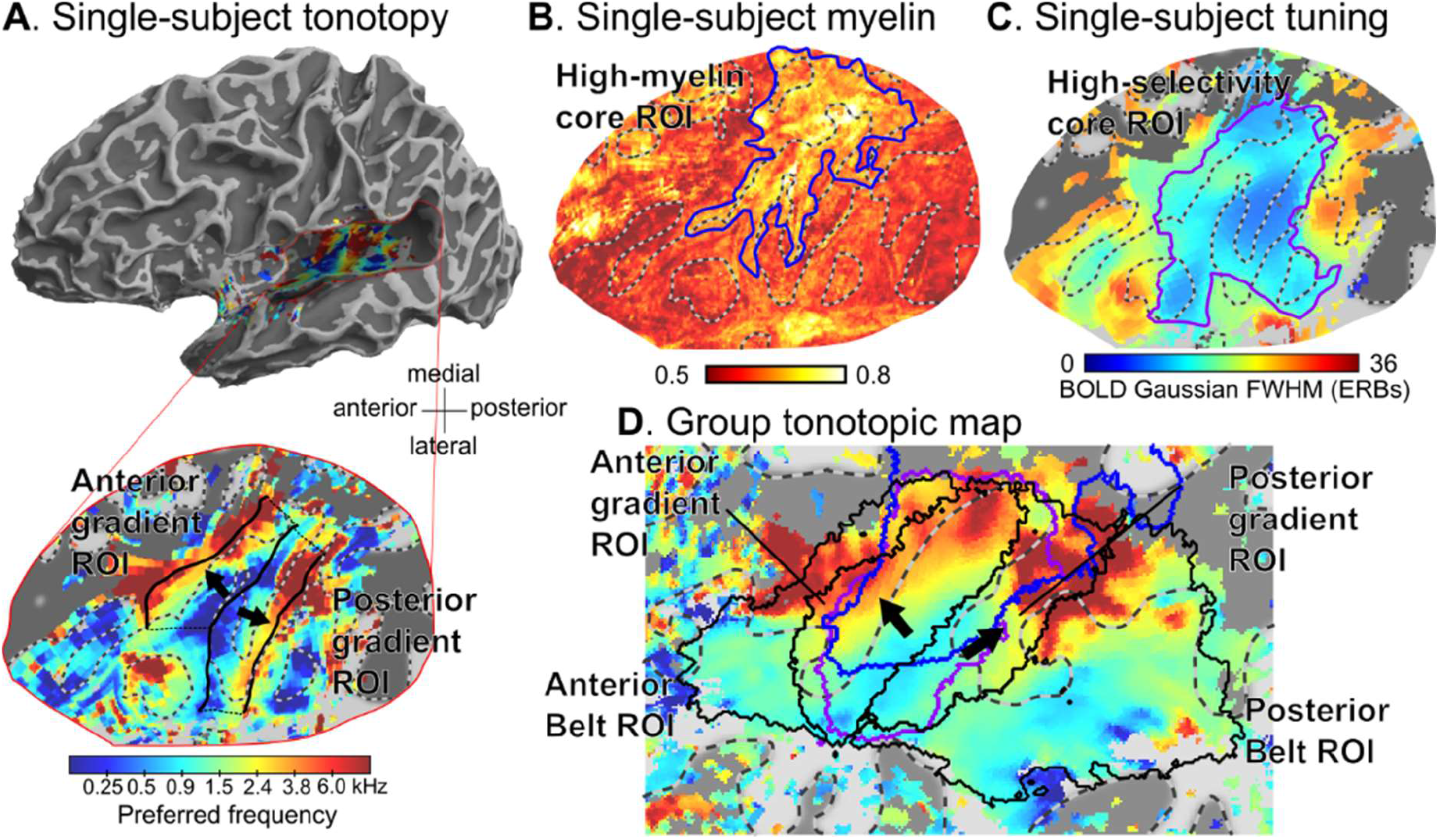
ROI definition at individual participant level and resulting probabilistic ROI location on group template. A. Map of voxelwise preferred-frequency map on semi-inflated (top row) and flattened (bottom row) cortical surface reconstructions of the left hemisphere and auditory cortex for participant 2. Preferred frequencies are shown on a quasi-logarithmic color scale (representing the cochlear frequency, or ERB-number, scale, see Material and Methods). Thick black solid lines show automatically-detected reversals in tonotopic gradients that were used to define the anterior and posterior gradient ROIs. Thin dotted lines join the reversals between the 2 gradient ROIs. Black arrows indicated the average direction of the tonotopic gradient within each ROI. Gyri are indicated by light-gray, and sulci by dark-gray highlight. The borders between them are shown by two-tone gray dashed lines. B. Individual map of PSIR-derived intracortical myelin content, proportional to the longitudinal relaxation rate R1. The blue outline show a region of high myelination, thresholded an individualized criterion [7]. C. Individual map of voxelwise BOLD response tuning width, measured as FWHM in cochlear ERBs. The purple outline highlights a region of high frequency selectivity (low tuning width). D. Group-average preferred-frequency map obtained by spherical normalization of the 22 individual hemispheres. Black, blue and purple outlines show the average location of the individual ROIs shown in A-C across the 22 left and right hemispheres, based on the maximum probability value of each ROI at each voxel of the template cortical surface (see Methods for details).

We then used the A, P and AP responses to estimate the suppressive effect of a given adaptor on the response to the probe. For this, we first estimated the adapted probe response, P|A, by subtracting A from AP (for corresponding adaptor frequencies). We then computed the adaptation effect by subtracting the adapted probe response P|A from the unadapted response P. Figure 3A-E shows group-average maps of the adaptation effect for increasing adaptor frequencies (averaged across all 11 participants, irrespective of the type of adaptor, single- or multiple-onset). There were significant adaptation effects for all adaptor frequencies but the amount of adaptation clearly increased as the adaptor frequency approached the probe frequency, suggesting that adaptation was frequency-specific. When the adaptor and probe frequencies were similar or equal (Fig. 3C-E), the adaptation effect was largest at cortical locations corresponding to the two high-frequency tonotopic gradient reversals separating each of the two gradient ROIs from the adjoining belt ROI (and presumably corresponding to the greatest response to the relatively high-frequency probe). In the posterior and, to a lesser degree, anterior belt ROI, even the lowest adaptor frequency, which was separated from the probe frequency by nearly three octaves, created significant adaptation. Finally, whilst the absolute adaptation amount was greatest in high-frequency-preferring regions (where the probe response itself was also greatest), the relative adaption amount (i.e., adaptation expressed as a proportion of the unadapted probe response, (P-A|P)/P) was similar across all regions, particularly, when the adaptor and probe frequencies were equal (Fig. 3F).

**Figure 3:**
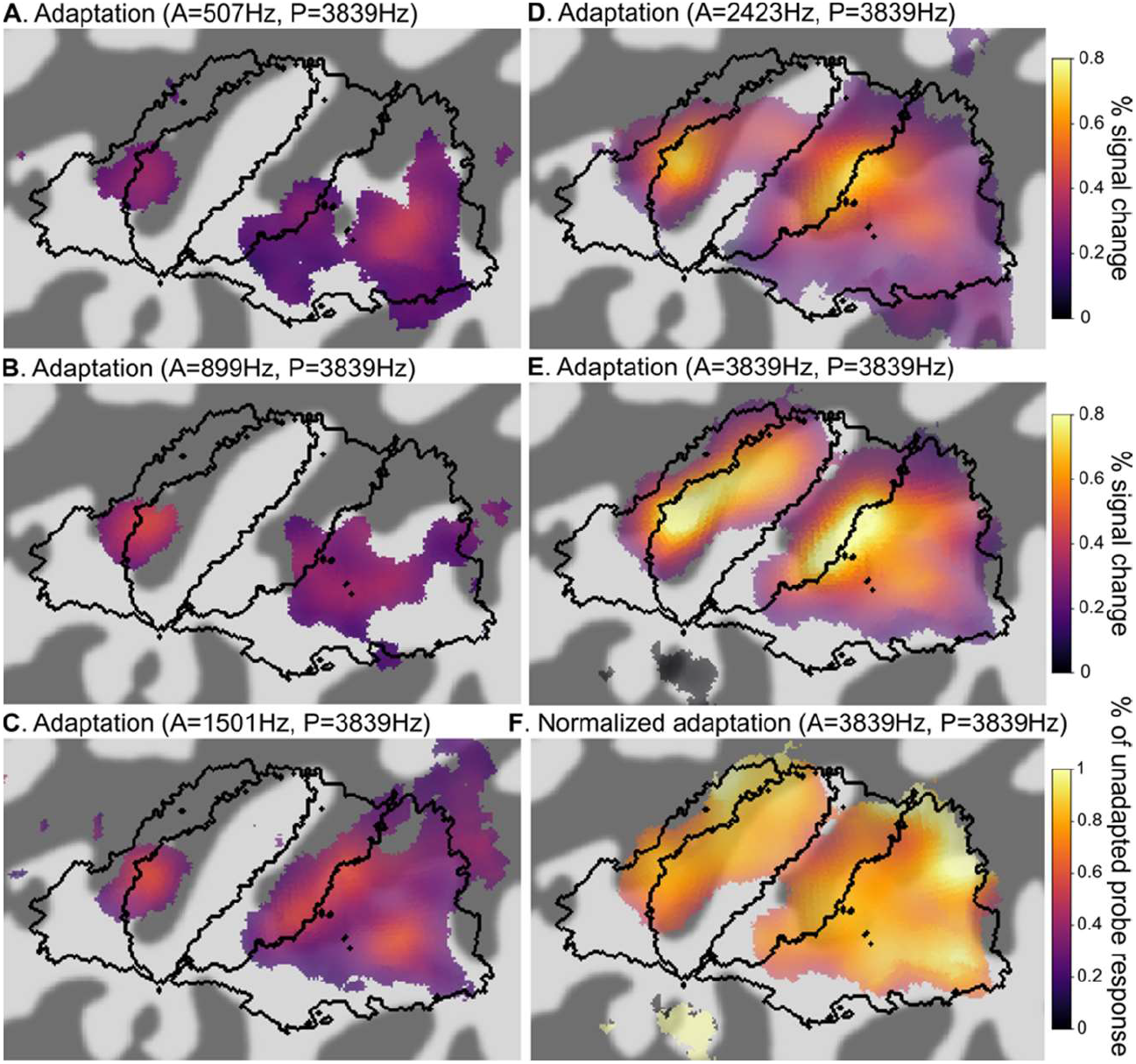
Group-level maps of the frequency adaptation effect for different adaptor-probe frequency differences (averaged across single- and multiple-onset adaptor conditions). A-E. Amplitude of the adaptation effect [P – P|A = P – (AP–A) = A + P - AP] for adaptors ranging between 0.51 and 3.84kHz and a 3.84kHz probe. Black outlines show the approximate location of ROIs (based on maximum probabilities, see Fig. 2D). Maps are, smoothed along the cortical surface (FWHM=6mm), thresholded at p<0.05 FDR-corrected and transparency is a power function of –log10(p). F. Same as E, but voxelwise adaptation values have been normalized by the corresponding voxelwise unadapted probe response [(P – P|A)/P]. All maps represent the average of the 22 left and right individual hemispheres.

### FMRI adaptation in auditory cortex is frequency-tuned following multiple-onset, but not single-onset, adaptors

To investigate the degree of frequency specificity of adaptation for each type of adaptor, we constructed adaptation tuning curves in the four ROIs by plotting the ROI-average adaptation effect as a function of the adaptor frequency (expressed in human cochlear frequency units or ERB-numbers). Figure 4A compares the adapted (P|A) to the unadapted (P) probe responses and Figure 4B plots the normalized adaptation tuning curves [(P-A|P)/P], separately for participants in the single- and multiple-onset conditions (light gray and black circles respectively). We found that the temporal properties of the adaptor (single-onset vs multiple-onset) led to markedly different adaptation tuning curves. Single-onset adaptors suppressed the probe response by similar amounts (∼40 to 60 %) irrespective of adaptor frequency, which resulted in a poorly-tuned adaptation effect. For multiple-onset adaptors on the other hand, the adaptation effect was clearly tuned in all four ROIs: suppression was small for low-frequency adaptors, and increased to a maximum (ranging from 74% in the anterior gradient ROI to 87% in the anterior belt ROI) when the adaptor frequency matched the probe frequency.

**Figure 4:**
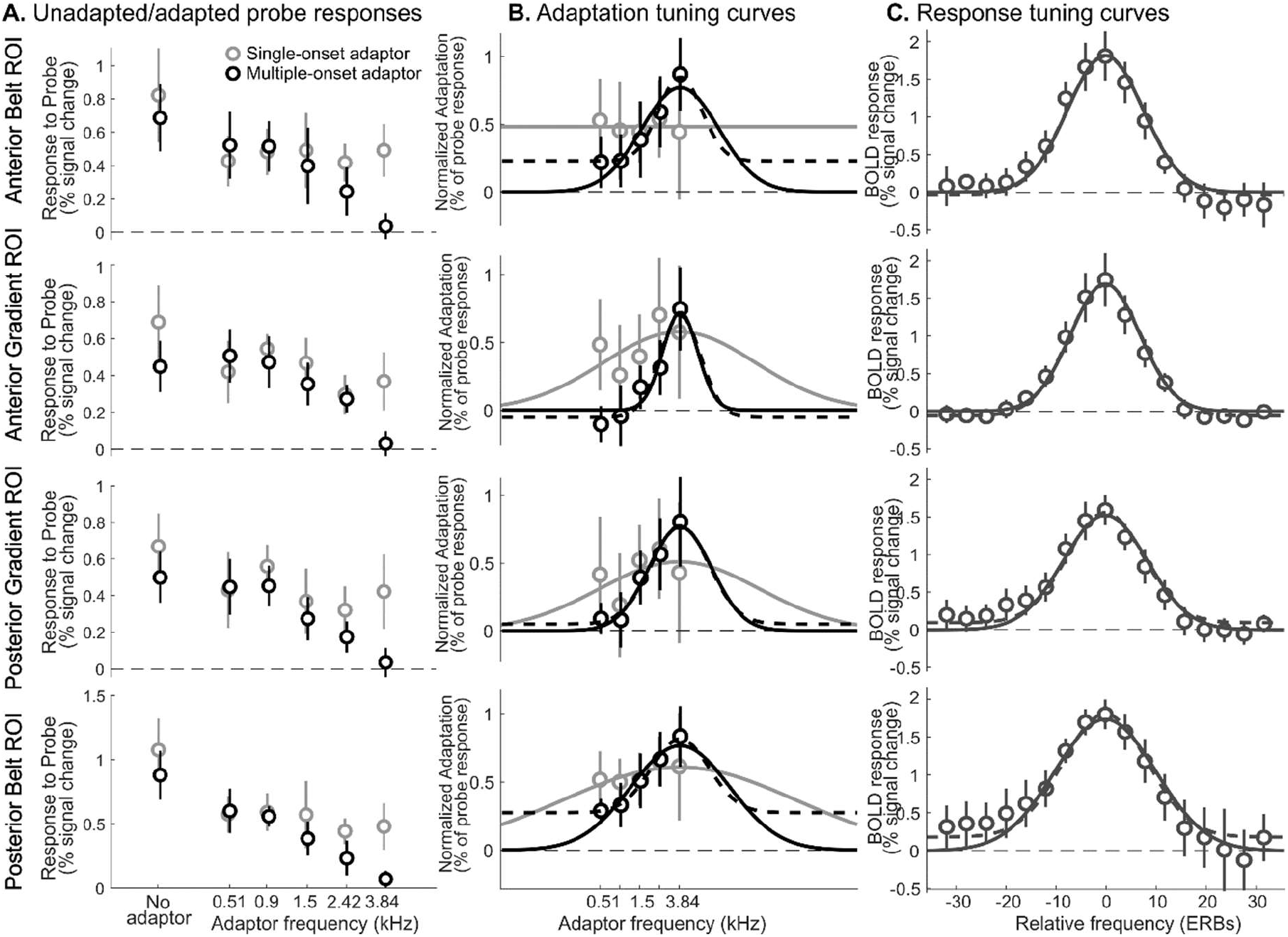
fMRI adaption and BOLD response tuning curves in core and belt region of auditory cortex. A. Unadapted (P) and estimated adapted (AP-A) BOLD responses to the probe in the two gradient (≈core) and the two belt ROIs, averaged across hemispheres for the five participants who were presented with a single-onset 200-ms adaptor (light gray circles) and for the six participants presented with multiple 50-ms adaptors (black circles). Adaptor frequencies on the X axis are equally spaced on the cochlear ERB-number scale, but expressed in kHz. B. ROI-average fMRI adaptation effect expressed as percent decrease from the unadapted response to the probe [(P – P|A)/P] as a function of the adaptor’s center frequency (i.e. fMRI adaptation tuning curve). The solid and dashed curves are the best-fitting Gaussians without and with a frequency-independent adaptation offset respectively (see Fig. 5 and supplementary Table S1 for best-fitting parameters). C. ROI-average voxelwise BOLD response to (multiple-onset) adaptors alone (A), as a function of the frequency difference between the A stimulus and the preferred frequency of each voxel (i.e. BOLD response tuning curve). Frequency differences on the X axis are expressed in ERBs. Solid and dashed curves as in panel B. In all panels, error bars represent 95% bootstrap confidence intervals.

To estimate the tuning widths of the adaptation effect for single- and multiple-onset adaptors, we fitted the adaptation tuning curves with Gaussian functions (solid curves in Fig. 4B; group-average best-fitting parameters and goodness-of-fit indicators given in supplementary Table S1). Simple Gaussian functions fitted the multiple-onset adaptation tuning curves reasonably well (average individual-hemisphere r^2^ ranging between 27 and 43% across ROIs), but provided a poor fit in the single-onset condition (r^2^ ranging between 0 and 2%). To test the degree of tuning in each condition, we compared the model evidence for the frequency-tuned Gaussian adaptation model (as quantified by either the Bayes Information Criterion, BIC, or the Akaike Information Criterion, AICc) against the evidence for a frequency-independent adaptation model in which adaptation suppression was constant across adaptor frequencies (i.e. equal to the average adaptation across frequencies). Both BIC and AICc provided weak-to-positive evidence against frequency-tuned adaptation in the single-onset condition, but very strong evidence for frequency-tuned adaptation in the multiple-onset adaptor condition (see supplementary Table S2). Since the remainder of this article is concerned with quantitative estimation of neuronal tuning width and single-onset adaptors did not result in frequency-tuned adaptation, all further analyses were conducted exclusively on the multiple-onset adaptor data.

Figure 5 plots the tuning width of the adaptation effect for the multiple-onset condition in each of the four ROIs, estimated as the FWHM of the best-fitting Gaussians. Adaptation tuning was narrowest in the anterior gradient ROI (8.2 ERBs, or ∼1.4 oct), wider in the posterior gradient ROI (14.8 ERBs, or ∼2.4 oct) and widest in the anterior and posterior belt ROIs (18.2 and 23.3 ERBs, or ∼3/3.8 oct; main effect of ROI: F(3,44) = 6.9184, p = .0006; pairwise comparisons only significant between the posterior belt ROI and each of the two gradient ROIs, p < .02).

**Figure 5:**
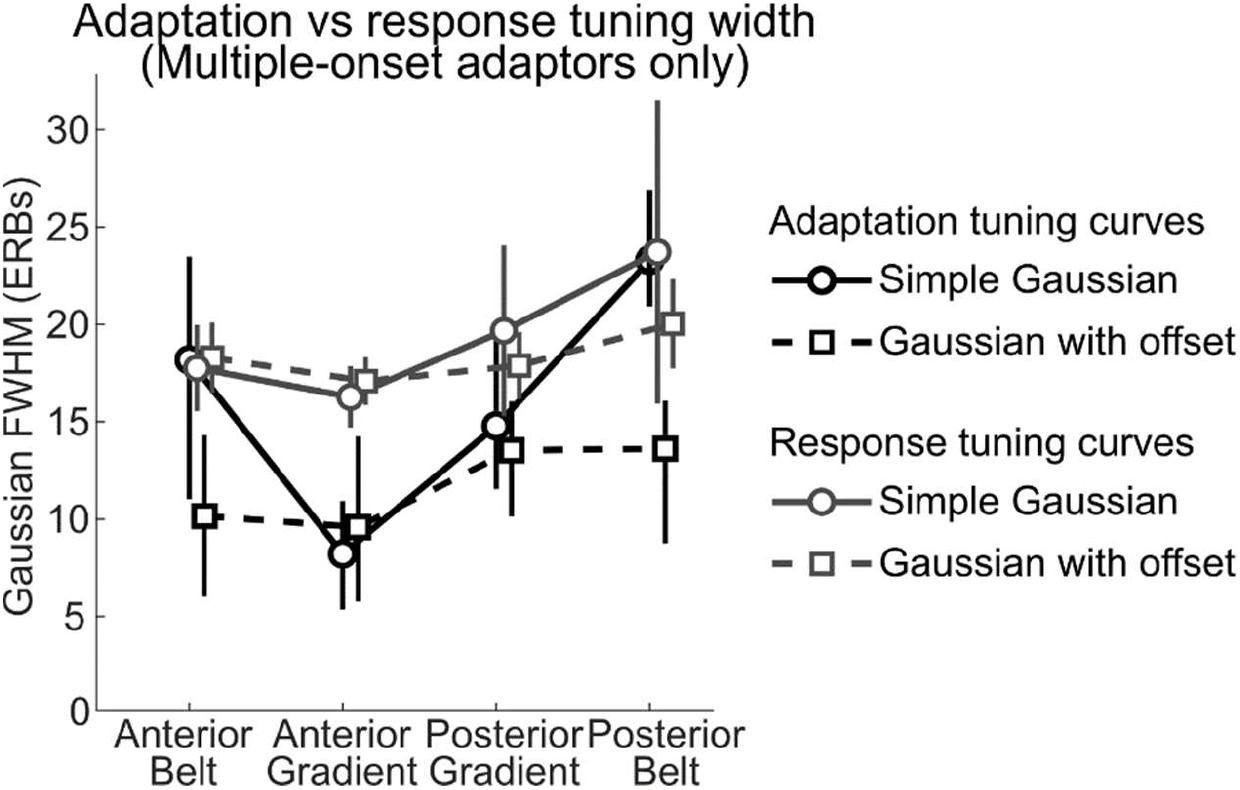
Estimated tuning width of fMRI adaptation and BOLD response tuning curves in gradient (core) and belt ROIs (black = adaptation; dark gray = response) for the six participants in the multiple-onset condition. Tuning width was estimated by fitting Gaussians to the group-average tuning curves, with or without frequency-independent adaptation offsets (dashed and solid traces respectively). Error bars represent 95% bootstrap confidence intervals. See supplementary Table S1 for average best-fitting Gaussian parameters obtained from individual hemisphere tuning curves.

Even though adaptation was clearly tuned in the multiple-onset adaptor condition, the minimum adaptation effect -when the adaptor-probe frequency separation was greatest – did not reach zero in all ROIs (see Fig. 4B): while it was virtually zero in both gradient ROIs (where adaptation was narrowly tuned), it was still substantial, around 25%, in both belt ROIs. This could be either because adaptation tuning curves in these regions were wider than could be measured using our finite range of adaptor frequencies, or because there was a residual frequency-independent baseline adaptation. Fitting simple Gaussians to estimate tuning width assumes that adaptation is entirely frequency-specific, and thus that the adaptation effect would approach zero for infinitely large adaptor-probe frequency separations. By not considering a possible frequency-independent adaptation component, this first fit may, however, have overestimated the tuning widths of the frequency-specific component, particularly in the belt ROIs. Hence, we fitted a second Gaussian model that included a constant offset, i.e. assuming instead that adaptation could reach a non-zero asymptote for infinite adaptor-probe frequency differences (dashed lines in Fig. 4B; average individual best-fitting parameters in supplementary Table S1). The best-fitting asymptotic adaptation values only differed from zero in the two belt ROIs, (posterior belt: 26%, p = .0002; anterior belt: 20%, p = .004). As a result, this second fit yielded substantially narrower adaptation tuning width estimates for the belt ROIs (dashed black lines in Fig. 5), with tuning widths ranging between 10.0 and 12.6 ERBs across all four ROIs, and no significant differences across ROIs (F(3,32.2) = 0.96, p = .42). Figure 4B suggests that the second fit yielded a better match to the data than the first fit for belt ROIs (also shown by larger r-squared values; see supplementary Table S1). However, this improvement in model fit did not lead to an increase in model evidence quantified through either the BIC or the AICc, which instead weakly favored the fit without asymptotic offset (see supplementary Table S2). This may be due to the large variance in adaptation values between participants and the limited range of adaptor frequencies used in our experiment.

In addition to these ROI Gaussian fits, we also fitted both types of Gaussian models to all voxelwise adaptation tuning curves. The resulting maps of voxelwise (or searchlight-) Gaussian parameters are shown in supplementary Figure S1.

### Comparison between fMRI adaptation and BOLD response tuning curves

Under the assumption that fMRI adaptation reflects underlying neuronal tuning properties, fMRI adaptation tuning would be expected to be narrower than the tuning of BOLD responses to different frequencies (see Introduction). To test this, we compared the tuning width of the fMRI adaptation effect to the tuning width of the BOLD response to adaptors presented alone. This analysis, as well as the remainder of the results, is based exclusively on the multiple-onset adaptor data because single-onset adaptors yielded poorly tuned adaptation effects.

To estimate the tuning width of the BOLD response, we constructed ROI-average voxelwise BOLD response tuning curves by averaging the voxelwise A response tuning curves, centered on the respective voxel preferred frequencies (i.e., the frequencies causing the maximum response in a given voxel; see Methods). The ROI-average voxelwise response tuning curves were approximately Gaussian-shaped when frequency was expressed in cochlear ERB-numbers (Fig. 4C), although they were slightly asymmetric, with larger responses at frequencies below than above the preferred frequency (except in the anterior gradient ROI). To measure response tuning width, we fitted these tuning curves with Gaussian functions (both with and without an asymptotic offset; Fig. 4C). Only the posterior belt ROI showed a small and (marginally) statistically significant response offset (p = 0.07; see Fig. 4C) and the response tuning width estimates (gray lines in Fig. 5) from the two fits were generally similar. They were smallest in the anterior gradient ROI (16.7/17.2 ERBs w/wo offset, corresponding to ∼3.1 octaves), and increased mainly towards the posterior belt region (fit without offset: F(3,30.78)= 6.57, p = .001, pairwise comparisons only significant between the posterior belt ROI and each of the two anterior ROIs, p < .02; fit with offset: F(3,33) = 13.566, p = 0.02, pairwise comparisons only significant between the posterior belt and anterior gradient ROI, p = 0.01).

Whether adaptation tuning width was smaller than response tuning width, and thus consistent with the expectation that fMRI adaptation reflects neuronal tuning properties, depended both on the type of Gaussian fit and on the ROI (Fig. 5): for the Gaussian fits without asymptotic offset, only the gradient ROIs showed narrower adaptation than response tuning, (anterior gradient ROI: F(1,22) = 10.52, p = .004; posterior gradient ROI: F(1,22) = 7.59, p = 0.02; belt ROIs: p > .82; note, however, that the interaction between ROI and tuning curve type did not reach significance: F(3,32.82) = 1.72, p = 0.18). In contrast, for the Gaussian fits with offset, adaptation tuning widths were consistently smaller than response tuning widths across all ROIs (main effect of tuning width type: F(1,11.11) = 43.81, p = .00004; comparisons between response and adaptation tuning widths significant at p < 0.02 in all ROIs; interaction between ROI and tuning curve type: F(3,65.27) = 1.13, p = 0.34).

### Using a computational model to explore the quantitative relationships between fMRI-adaptation, BOLD-response and neuronal tuning

Our results so far suggest that, at least within the core region (gradient ROIs) and when using trains of multiple adaptors, fMRI adaptation tuning is sharper than BOLD response tuning – consistent with the idea that fMRI adaptation tuning can reflect neuronal tuning. To better understand how adaptation and response tuning quantitatively relate to the underlying neuronal tuning properties, we devised a simple computational model of fMRI adaption in a unidimensional cortical strip of frequency-tuned neurons (referred to as tonotopic array; illustrated in Fig. 6A,E). In this model, we assume for simplicity that the adaptation tuning width of a neuron directly reflects its output response tuning and is therefore not inherited from the neuron’s input. We show that, even under this assumption, fMRI adaptation overestimates neuronal tuning width.

**Figure 6:**
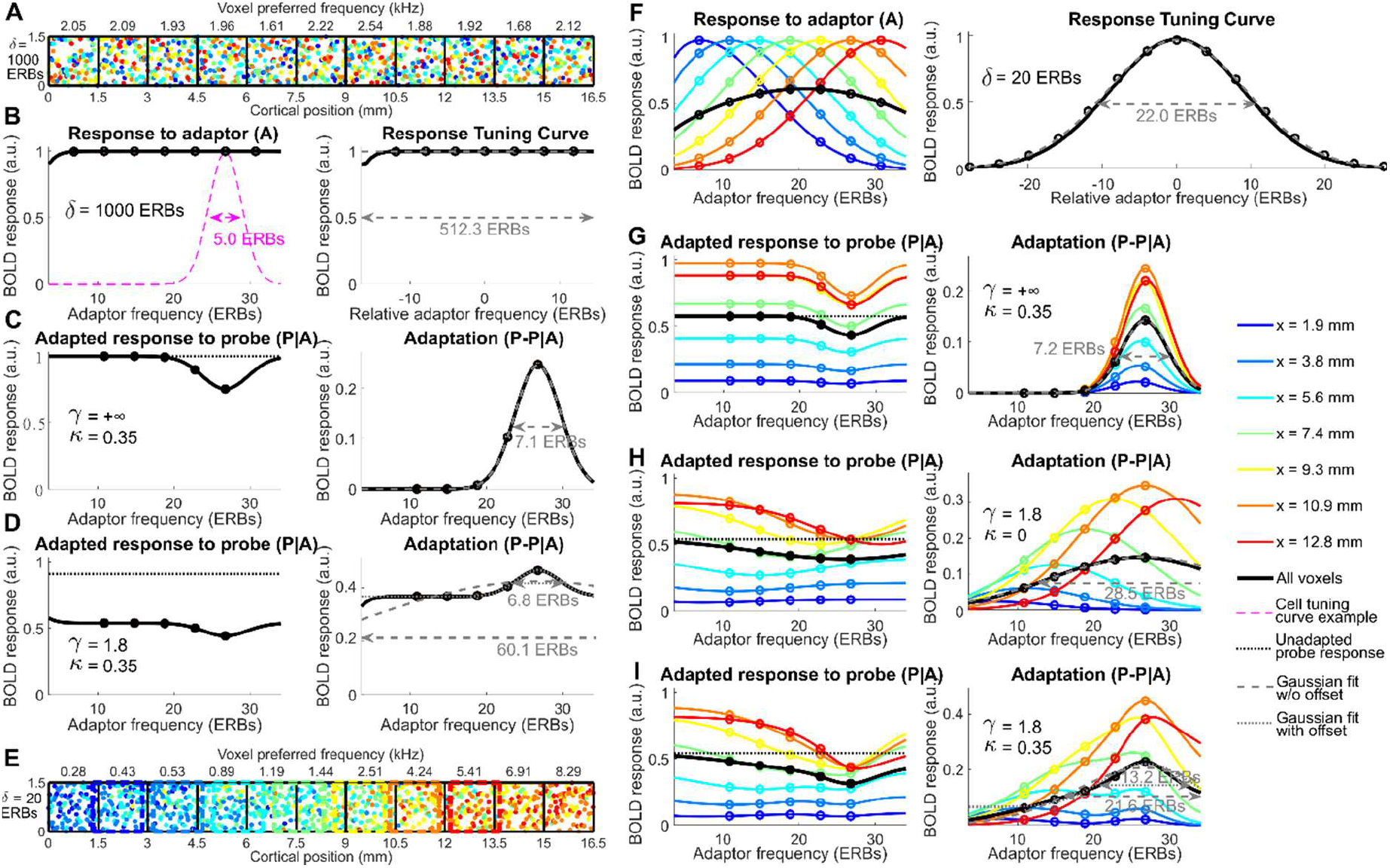
Simulated BOLD response and BOLD adaption tuning curves in a cortical model of fMRI adaptation for different tonotopic scatter, neuronal adaptation strength and BOLD saturation parameters. A. Graphical illustration of the spatial layout of the model without tonotopic organization (tonotopic scatter δ = 1000 ERBs). The black grid represents contiguous voxels extending the full length of the model and the colored dots represent the spatial cortical position of 100 randomly-drawn neurons per voxel, with the color of the dot coding for best frequency (blue = low frequency and red = high frequency). The actual model also included intermediate voxels overlapping with the represented voxels. Figures above each voxel give the estimated preferred frequency of the voxel. Although neurons and voxels are represented here in 2D for clarity, the actual model was unidimensional (the represented transverse dimension was collapsed). Because of the lack of tonotopic organization, the modelled BOLD response was identical at all of voxels and is illustrated in panels B to D. B. Modelled voxel BOLD responses to adaptor-only trials (A) as a function of adaptor frequency (left panel) and BOLD response tuning curve (right panel) obtained by averaging voxel tuning curves re-centered on their respective center frequencies. In both panels, open dots correspond to the seven adaptor frequencies actually used in our fMRI experiment, while solid traces show responses for a more extensive range of adaptor frequencies between 0.19 and 7.82 kHz). In the left panel, the response to the 3.84 kHz adaptor also corresponds to the unadapted response to the probe (P). The dashed magenta trace illustrates the tuning curve of an auditory neuron responding maximally to the probe with a tuning width of 5 ERBs FWHM. The modelled BOLD response tuning curve (right panel) was fitted with a Gaussian (gray dashed line) whose FWHM (gray double-arrow) is reported in ERBs. C. Adapted BOLD response to the probe as a function of adaptor frequency (left panel) and unnormalized fMRI adaptation tuning curve (right panel) computed by subtracting the adapted probe response from the unadapted probe response (replotted from the left panel of B as a horizontal dotted line), for an instance of the model including neuronal adaptation (neuronal adaptation strength parameter κ = 0.35) but no neurovascular compression (BOLD saturation parameter γ = +∞). D. Same as C, but the BOLD saturation γ was set to 1.8. The fMRI adaptation tuning curve (right panel) was fitted both with a simple Gaussian (gray dashed line) and a Gaussian with an offset parameter (gray dotted line). Corresponding Gaussian FWHM are reported below the double arrows. E. Same as A, but the tonotopic scatter was reduced to δ = 20 ERBs, resulting in a tonotopically-organized model. The modelled BOLD responses of the seven voxels outlined with colored dashed lines are illustrated in panels F to I in the corresponding color. F. Same as B with δ = 20 ERBs instead of 1000 ERBs. G. Same as C, but with δ = 20 ERBs instead of 1000 ERBs. H. Same as G, but with neurovascular compression (γ = 1.8 as in D) and without neuronal adaptation (κ = 0). I. Same as G and H, but with both neuronal adaption (κ = 0.35) and neurovascular compression (γ = 1.8).

In the model, neuronal tuning curves were assumed to be Gaussian-shaped on the ERB-number scale (dashed magenta curve in Fig. 6B) with nominal characteristic frequencies increasing from one side of the array to the other. Neuronal tuning width, *κ*, was assumed to be constant and identical for all neurons. While the nominal characteristic frequency of neurons increased tonotopically, their actual characteristic frequency at a given location within the array was allowed to vary according to a Gaussian distribution of width δ, which controlled the degree of tonotopic scatter. Parameters *κ* and δ will be expressed as Gaussian FWHM throughout the manuscript. The model implemented adaptation both at the neuronal and BOLD levels [the latter reflecting non-linearities in neurovascular coupling, see e.g. 28]. Neuronal adaptation was implemented as a simple multiplicative suppression, with the suppression amount proportional to the size of the neuron’s adaptor response (i.e., assuming a response fatigue adaptation mechanism, see Discussion). The maximum amount of neuronal adaptation (when the adaptor and probe frequencies were equal) was controlled by a proportionality parameter, *κ*, henceforth referred to as neuronal adaptation strength. Neurovascular non-linearity was implemented through a compressive sigmoid function, which was applied to the aggregate BOLD response of all neurons contained within a given voxel. The saturation parameter γ controlled the maximum possible response and thus the degree of compression (see Methods section for more implementation details).

To informally test the behavior of the model and its ability to reproduce our experimental results, we first set the model’s free parameters (*σ, δ, κ* and γ) manually. Across all informal tests, we set the neuronal tuning parameter, *σ*, to 5 ERBs (shown in Fig. 6B) – close to the average cellular tuning width reported for non-human primates (see Discussion).

We first tested a version of the model without tonotopic organization (by setting tonotopic scatter, *δ*, to a large number – 1000 ERBs). In this version, neuronal characteristic frequencies were essentially uniformly distributed across the array (Fig. 6A), and thus BOLD response tuning curves (response to A-only stimuli) were flat (Fig. 6B). This was true for both the voxelwise responses and the average responses across all voxels within the array, irrespective of whether or not the voxelwise tuning curves were re-centered on their preferred frequencies before averaging (compare left and right panels in Fig. 6B). Figure 6C (left panel) shows voxelwise and array-average adapted probe responses for different adaptor frequencies when there was no neurovascular compression (γ = +∞), and thus, only neuronal adaptation was at play (*κ* = 0.35). In this scenario, the probe response was only suppressed for adaptor frequencies close to the probe frequency, resulting in a narrowly tuned adaptation tuning curve (Fig. 6C, right panel). The estimated adaptation tuning width (measured in FWHM) was 7.1 ERBs, which is a factor of 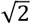larger than the underlying 5-ERB neuronal tuning width. In this non-tonotopic model, introducing neurovascular compression (by setting γ to 1.8) resulted in an additional, general reduction of the adapted probe responses (Fig. 6D, left panel) and a corresponding frequency-independent adaptation component in the array-average adaptation tuning curve (Fig. 6D, right panel). Fitting this adaptation tuning curve with a simple Gaussian (i.e., without asymptotic offset) yielded a tuning width of 60.1 ERBs – more than an order of magnitude larger than the underlying neuronal tuning width (5 ERB). However, fitting a Gaussian with offset yielded an adaptation tuning width of 6.8 ERBs – similar to the estimated adaptation tuning width in the absence of neurovascular compression (7.1 ERBs).

The above simulations demonstrate that, for neuronal populations that are not topographically organized, the stimulus-specific component of fMRI adaptation tuning is directly related to underlying neuronal tuning, but overestimates neuronal tuning width by a factor 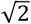. Neurovascular compression also contributes to the suppression of the probe response, but this effect is stimulus-independent and can thus easily be separated from the stimulus-specific suppression caused by neuronal adaptation. We will now show that this simple distinction between neurovascular and neuronal adaptation effects ceases to apply when the degree of tonotopic order increases to a point where response tuning becomes apparent at the voxel level. To demonstrate the effect of BOLD response tuning, we set the degree of tonotopic scatter, *δ*, to 20 ERBs. With this lower degree of scatter, voxelwise average neuronal characteristic frequencies increased systematically across voxels (Fig. 6E) and, as a result, the voxelwise response tuning curves (Fig. 6F, left panel) became peaked, with maximum responses near the average neuronal characteristic frequencies. The resulting recentered, array-average response tuning curve (Fig. 6F, right panel) was approximately Gaussian-shaped with a width of 22.0 ERBs, similar to the response tuning width actually measured in the posterior belt ROI (parameter *δ* was fine-tuned to 20 ERBs to achieve this similarity). When we assumed that the only source of adaptation was neuronal (*κ* = 0.35 and γ = +∞), the adapted probe responses again showed a localized trough around the probe frequency (Fig. 6G, left panel), and the array-average adaptation tuning curve showed a corresponding peak (Fig. 6G, right panel), despite the unadapted probe response varying widely in amplitude across voxels. The width of the array-average adaptation tuning curve was again 7.2 ERBs, approximately 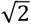times the underlying neuronal tuning width, as for the non-topographic model with no neurovascular compression (compare with Fig. 6C, right panel). When, on the other hand, we assumed that the only source of adaptation was neurovascular compression (by setting *κ* to zero and γ to 1.8), the voxelwise adaptation tuning curves became broadly tuned (Fig. 6H), with peak locations corresponding to each voxel’s preferred frequency, rather than the probe frequency (compare with Fig. 6F). This is because suppression due to neurovascular compression increases with increasing adaptor response and is therefore strongest when the adaptor frequency matches the voxel preferred frequencies. As the unadapted probe response size varies between voxels depending on their tuning properties, different voxels contribute differently to the array-average adaptation effect (Fig 6H, right panel), with voxels preferring frequencies close to the probe frequency (3.84 kHz) contributing more than voxels preferring lower frequencies. Thus, the array-average adaptation tuning curve appeared to be tuned to the probe frequency, but with a much broader tuning width than in the model with purely neuronal adaptation (28.5 ERBs; compared black curves in Fig. 6G and H, right panels). Finally, when assuming both neuronal adaptation and neurovascular compression as sources of fMRI adaptation, the voxelwise and array-average adaptation tuning curves (Fig. 6I) showed a superposition of the broader and more narrowly tuned adaptation effects seen in the previous two models (Fig. 6I, right panel). Fitting the resulting array-average adaptation tuning curve with a simple Gaussian (Fig. 6I, right panel) yielded an intermediate adaptation tuning width of 21.6 ERBs, which, as for the measured data in belt regions, was similar to the width of the array-average response tuning curve (22.0 ERBs; compare with Fig. 6F). Fitting with a Gaussian with constant offset, on the other hand, yielded a narrower adaptation tuning width (13.2 ERBs), again similar to the measured data (compare with Fig 4B and Fig. 5, dashed black line), but still wider than the tuning width of the purely neuronal fMRI adaptation effect (7.1 ERBs) and corresponding underlying neuronal tuning (5 ERBs).

The above informal simulations suggest that fMRI adaptation not only overestimates the underlying neuronal tuning by a systematic 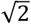factor, but also that, in topographically-organized cortex, neurovascular non-linearity manifests as a frequency-specific adaptation effect that is difficult to disentangle from effect of neuronal adaptation, leading to further overestimation of neuronal tuning width. In Supplementary Figure S2, we show that these biases exist for a wide range of model parameters.

### Can cortical single-neuron frequency selectivity be inferred from fMRI adaptation data?

In this section, we test whether we can use our fMRI adaptation model to disentangle neurovascular and neuronal adaptation effects in our fMRI adaptation data and thereby obtain a direct estimate of average single-neuron tuning widths in different auditory regions. To achieve this, we fitted the model to the observed group-average BOLD response and fMRI adaptation tuning curves across the fours ROIs. Three of the model’s free parameters – the tonotopic scatter, *δ*, the neuronal tuning, *σ*, and neuronal adaption strength, *κ* – were assumed to vary across ROIs, while the BOLD saturation parameter, γ, was assumed to be constant across ROIs. We fitted the model 1000 times with random initial parameters. Figure 7A,B shows the median fitted response and adaptation tuning curves compared to the initial curves before fitting. The fitted curves (thick black traces) matched the measured data (dark gray symbols) considerably better than the initial curves (thin light gray traces), with average fitted vs initial r^2^ of 70.0% vs 15.6%. Even the fitted curves, however, did not capture all features of the measured data. Specifically, they did not reproduce the asymmetric offset of the response tuning curves observed in most ROIs (Fig. 7A) – as expected, since tuning is entirely symmetric in the model. They also did not generally fit the adaptation tuning curves as well as the previous Gaussian models (Fig. 7C), especially in the gradient ROIs. In the belt ROIs, the fitted curves approximately followed the simple Gaussian fits, but did not follow the apparent frequency-independent adaptation component captured by the Gaussians with offset.

**Figure 7:**
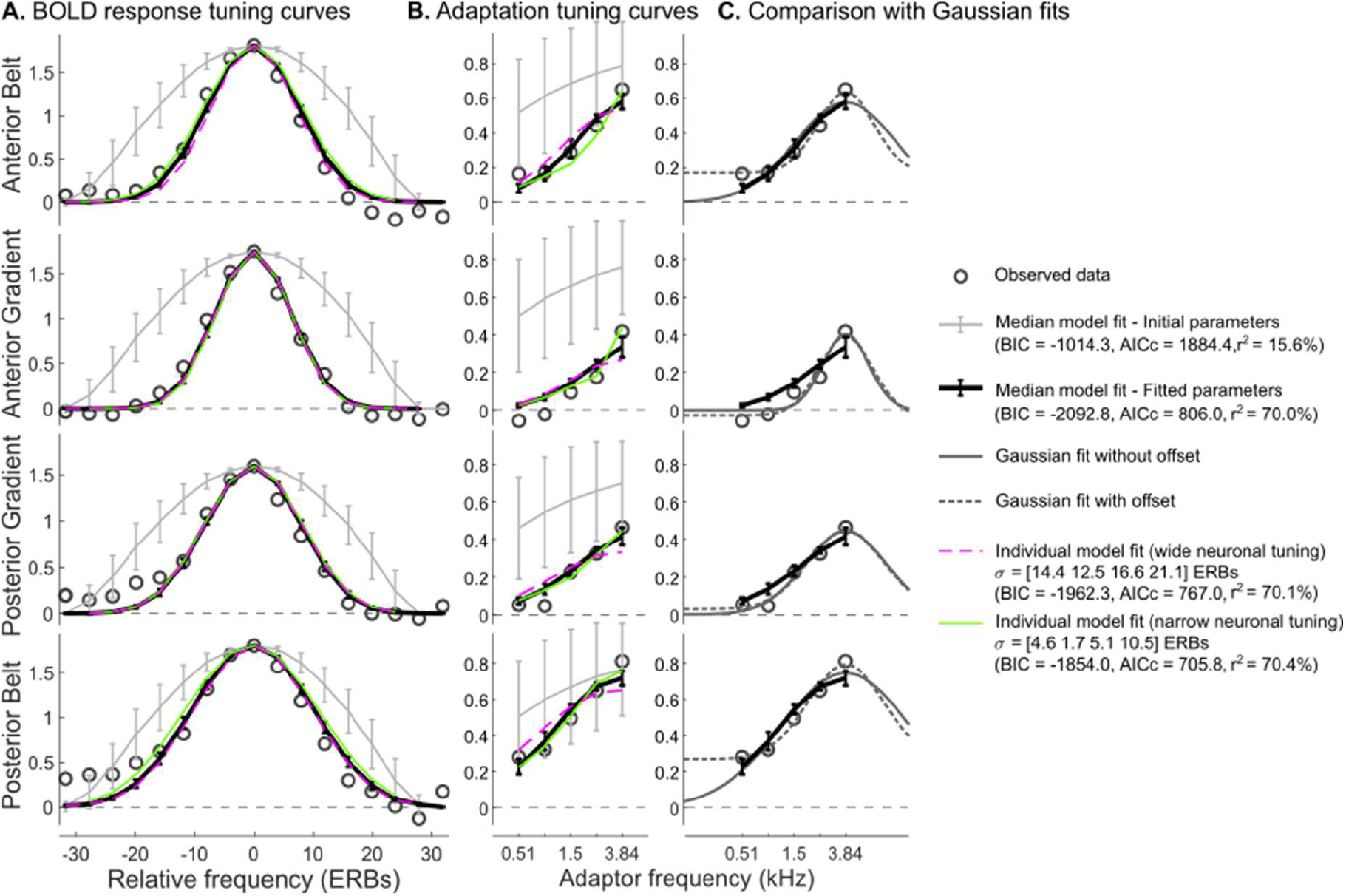
Modeled BOLD response (A) and fMRI adaption tuning curves (B) superimposed on the corresponding experimental ROI-average tuning curves. Dark gray open circles represent the actual tuning curves, replotted from Fig. 4B-C. Thick black traces represent the median of 1000 best-fitting model tuning curves and thin light gray traces the median of the correspond 1000 initial parameter tuning curves. Error bars are interquartile intervals across the 1000 fits. The coloured traces correspond to two example individual fits, with parameters highlighted in Fig. 8. The dash magenta trace corresponds to an individual fit with wide neuronal tuning parameters and the solid green trace to a fit with narrow neuronal tuning parameters. (C) Comparison between the actual and fitted adaptation tuning curves, replotted from panel B and the two types of Gaussian fits (with and without asymptotic offset) replotted from Fig. 4B,C.

Figure 8A shows the distributions of the individual fitted parameters for each of the 1000 fits (colored solid lines), compared to the (uniform) distribution of the random initial parameters (gray dashed line). Whilst the fitted distributions were narrower than the corresponding initial distributions, most of the fitted distributions were still relatively wide, with at least two pronounced peaks (with the exception of the neuronal adaptation strength, *κ*, whose distribution was unimodal). At the same time, all parameters showed marked distributional differences between ROIs. The anterior gradient ROI had the smallest neuronal tuning widths (*σ* = 0.2 to 14 ERBs), as well as the smallest degree of tonotopic scatter (*δ* = 1 to 15 ERBs) and the least amount of neuronal adaptation (*κ* = 0.4 to 0.65, at par with the posterior gradient ROI). The posterior belt ROI, on the other hand, had the corresponding highest values for these three parameters, and the posterior gradient and anterior belt ROIs showed intermediate (and similar) values for *σ* and *δ*. Notably, best-fitting neuronal adaptation strengths were lower in the two gradient ROIs than in the two belt ROIs. The degree of neurovascular compression (as represented by the reciprocal of the BOLD saturation parameter 1/γ), common to all four ROIs, ranged between 0.05 and 0.63 (corresponding to γ parameter values between 2.8 and 36% signal change when rescaled to the measured BOLD responses). Scatter plots between pairs of the fitted parameters (Fig. 8B) indicated covariance between them, explaining their wide and multi-peaked distributions. Neuronal tuning width, *σ*, and tonotopic scatter, *δ* (leftmost panel) were negatively related with what looked like a quadratic relationship (*σ*^2^ + *δ*^2^ = *constant*). On the other hand, *σ* was negatively related to neurovascular compression, 1/γ. These patterns of covariance suggest a continuum of model solutions between two extremes: a model with narrow neuronal tuning (< 10 ERBs in all ROIs), high degree of tonotopic scatter (12 to 25 ERBs, depending on the ROI) and strong neurovascular non-linearity (i.e., 1/γ between 0.4 and 0.66, corresponding to saturation parameter, γ, between 2.7 to 4.5% signal change), and an opposite model with relatively wide neuronal tuning, low tonotopic scatter, high neuronal adaptation strength and weaker neurovascular non-linearity (*σ* = 12 to 24 ERBs, *δ* < 10 ERBs, 1/γ < 0.4 or γ > 4.5 % signal change).

**Figure 8:**
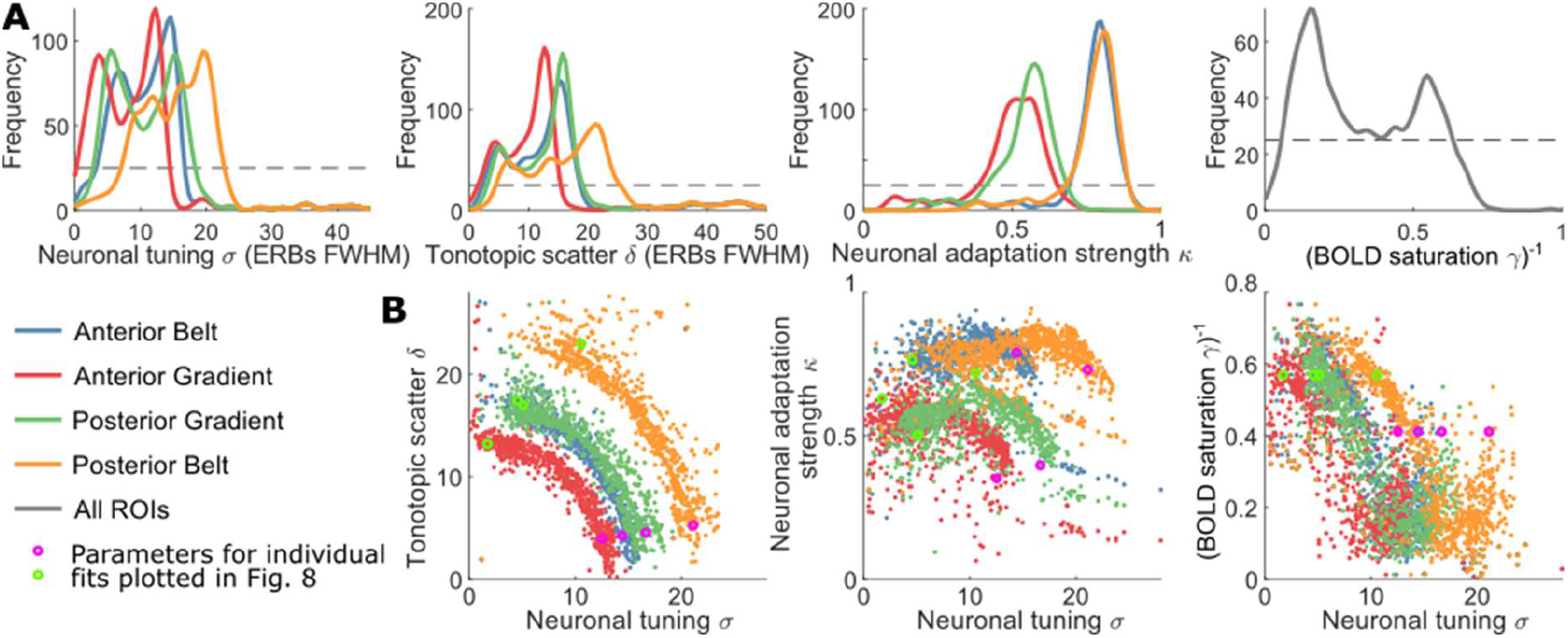
Distributions of best-fitting parameters across 1000 fits with random initial parameters.A. Distributions of the best-fitting parameters for the 4 free parameters in the 4 core and belt ROIs (= 10 free parameters since γ was common to all ROIs). Distributions were smoothed using kernel density estimation. The thin dashed black line represents the uniform distribution of random initial parameters. B. Selected joint distributions of pairs of free parameters across the 1000 fits, plotted as scatter plots for each of the 4 ROIs. Parameters sets corresponding to the two individual fits plotted in Fig. 7 are highlighted in magenta and green, respectively.

Two example model fits (one at each extreme of this continuum) are illustrated in Figure 7A,B (green and magenta dashed traces for narrow and wide neuronal tuning respectively, with circles of corresponding colors in Fig. 8B showing the respective parameters). Both fits were very similar and closely followed the median fitted model, with the exception that the model with narrow neuronal tuning (green trace) seemed slightly better at accounting for both the peak of the adaptation curves and the frequency-independent adaptation component (in the belt ROIs).

## Discussion

Using an fMRI adaptation paradigm, we demonstrated frequency-specific fMRI adaptation in human primary and secondary auditory cortex, such that the amount of suppression increased as the frequency of the adaptor approached that of the probe. Frequency-specific adaptation was only observed for short trains of multiple adaptors however, not for single-onset adaptors. For multiple-onset adaptors, fMRI adaptation frequency tuning curves were narrower than BOLD response tuning curves, at least in primary cortex, consistent with the assumption that fMRI adaptation can reflect neuronal frequency tuning. Using computational modelling, we identified two sources of bias that can cause population (fMRI) adaptation tuning to overestimate the underlying neuronal tuning. Directly fitting our model to the fMRI adaptation data is a first step towards estimating average neuronal frequency tuning in human core and belt auditory cortex non-invasively.

### Quantitative estimation of neuronal tuning width from fMRI adaptation

FMRI adaptation has been suggested to reflect neuronal selectivity [14, 15, 29], but to quantitatively infer single-neuron tuning curves from fMRI adaptation measured in a voxel or in an ROI, the following relationships must be accounted for: (1) how single neurons’ output (spike-based) tuning curve relates to their adaptation tuning curve, (2) how adaptation tuning at the level of a neuron relates to adaptation tuning of the level of the neuronal population that contributes to the fMRI adaptation effect and (3) how the BOLD response relates to this population’s neuronal response. Here we discuss each of these three relationships with respect to our fMRI and modelling results.

Although it might be naively assumed that adaptation tuning directly reflects output tuning in a given neuron, as we did in our model, and consistent with adaptation being exclusively due to spiking fatigue mechanisms, such as hyperpolarization [30], neuronal adaptation and response tuning curves have often been shown to differ, both in visual and auditory cortex [26, 31-34]. This difference has generally been interpreted as suggesting either that adaptation reflects additional adaptive synaptic mechanisms at the neurons’ input, such as synaptic depression [35-37], or that it is inherited from upstream regions that project to the recording site [23, 27]. In either case, the tuning of the adaptation effect measured in a given cortical region could reflect the tuning of neurons located in different (upstream) brain regions. To try and modulate the contribution of this inherited adaptation in our fMRI adaptation paradigm, we manipulated the temporal properties of the adaptor. Multiple successive adaptors have been shown to result in more frequency-specific adaptation effects than single-onset adaptors [38, 39], as we also observed here. This may be due to the fact that for repeated or prolonged presentation of the adaptor, adaptation of upstream neurons allows downstream neurons to recover from adaptation, and the adaptation effect therefore reflects the sharper tuning of upstream neurons [40]. According to this explanation, the tuning of the adaptation effect should be less influenced by inherited adaptation for single or short adaptors than for long or multiple adaptors. Here we did observe a difference in tuning in the predicted direction, but found that only multiple-onset adaptors resulted in clearly frequency-tuned adaptation. Therefore the adaptation tuning widths we report, which are based on the multiple-adaptor condition, may reflect the neuronal tuning of upstream brain regions. It is unlikely however that the observed fMRI adaptation effect exclusively originates from upstream subcortical structures, because we also found clear differences in adaptation tuning between cortical areas, suggesting that at least part of the adaptation effect is local to each region. It is also important to emphasize that cortical connections are predominantly local and recurrent, and therefore that upstream and downstream neurons are often located in the same region [14]. Note that, contrary to previous studies, we equated the total duration of single-onset and multiple-onset adaptor trains (200 ms). The fact that we still observe a difference in tuning between the two conditions suggests that stimuli with multiple onsets are more efficient adaptors and that onset response plays a special role in adaptation.

Independently of whether, or to what extent, adaptation tuning reflects neuronal tuning at the site of recording, our modelling results highlight two additional factors that must be taken into account in order to quantitatively derive neuronal tuning from fMRI adaptation. The first concerns the relationship between the tuning of a single neuron and the tuning of a population of neurons measured at the macroscopic level, which differed by a factor 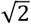 in our simulations. According to the logic of the adaptation paradigm [14, 15, 29], the tuning of adaptation to a given feature (sound frequency in our case) should reflect the tuning of a subpopulation of neurons (within a voxel or ROI) that are sensitive to this feature. For instance, in the adaptor-probe paradigm we used here, the fMRI adaptation tuning curve should reflect the tuning of neurons responsive to the probe. This is supported by our results, which showed the strongest adaptation effects in high-frequency-preferring regions, which also produced the strongest probe responses. This, however, does not mean that the aggregate adaptation tuning curve at the population level should exactly match the adaptation tuning of single neurons. Amongst neurons that respond to the probe frequency, and therefore contribute to the adaptation effect, some have a response tuning curve centred on the probe, but many others may prefer other frequencies, while still significantly responding to the probe. The aggregate fMRI adaptation tuning curve should therefore be wider than the adaption tuning curve of any individual neuron. In the supplementary discussion, we formally demonstrate that the contribution of off-frequency neurons should widen the population adaptation tuning curve by a factor 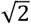compared to the underlying neuronal tuning width, consistent with our simulation results. This bias occurs in the absence of any other confounding factors (such as neurovascular non-linearity, discussed in the next paragraph) and would therefore be expected to exist in any macroscopic measurement of adaptation tuning curves, including

EEG adaptation. It also exists independently of whether the feature of interest is topographically organized in the probed region, and would therefore apply to previous fMRI adaptation that attempted to quantitatively measure neuronal tuning for orientation or motion direction in visual cortex [19-21].

The second factor is the contribution of neurovascular non-linearity to fMRI adaptation, specifically in topographically organized cortex. It is well known that the BOLD response to two stimuli presented in close succession is subadditive [e.g. 41]. Although a large part of this sub-additivity might be neuronal in origin and could therefore itself reflect stimulus-specific neuronal adaptation [42, 43], it likely also includes non-neuronal (e.g. neurovascular or vascular) contributions [28, 44, 45], which would not generally be expected to show stimulus specificity. We modelled this non-neuronal sub-additivity by applying an unspecific non-linearity (saturating function) to the voxel BOLD responses, and found that the effect of this non-linearity – despite itself being unspecific – could still appear stimulus-specific when the neuronal population was given a sufficient degree of topographic organization for the voxel BOLD responses to be frequency-tuned at the voxel level (as is the case for frequency preference in auditory cortex). This apparent stimulus specificity of the neurovascular non-linearity effect came about as a result of averaging the absolute adaptation amounts across voxels. As the absolute adaptation amounts depend on the sizes of the voxels’ unadapted probe responses, voxels tuned to frequencies close to the probe frequency contribute more strongly to the average adaptation effect than voxels tuned to more remote frequencies, resulting in apparent tuning. This neurovascular adaptation tuning should be wider than neuronal adaptation tuning because it reflects the tuning of the BOLD responses rather than of individual neurons, thus leading to additional overestimation of the neuronal response tuning width.

While neurovascular non-linearity may have contributed to the observed frequency-specific adaptation effect, it cannot explain the apparent frequency-independent adaptation component observed in the auditory belt regions, because voxels in these regions showed clearly frequency-tuned BOLD responses. This frequency-independent adaptation component could instead reflect the observed asymmetrical shape of the BOLD response tuning curve: the BOLD response tended to be above baseline for frequencies well below the voxel preferred frequencies, which may have translated into non-zero adaptation for adaptor frequencies well below the probe frequency. Asymmetric BOLD response tuning curves may reflect a similar asymmetry of cochlear tuning curves [4, 46], although, if this is the case, it remains unclear why we did not observe this asymmetry in the core region. Further measurements with a wider range of adaptor frequencies on both sides of the probe will be necessary to ascertain origin of the frequency-independent adaptation component.

### Comparison with previous cortical frequency tuning measurements

Fitting our fMRI adaptation model to the measured response and adaptation tuning curves allowed us to estimate average neuronal tuning properties in different auditory cortical regions [47, 48], while accounting for (at least some of) the measurement-related biases described in the previous section. Given the covariance between the free parameters of the model, we could only obtain a fairly wide range of estimates for neuronal frequency tuning, but these are consistent with neuronal tuning being narrower in core (0.1-14 ERBs in the most selective region in the anterior gradient ROI overlapping most with core auditory cortex on Heschl’s gyrus) than in belt auditory cortex (7-23 ERBs for the least tuned region in the posterior belt ROI on planum temporale), as has been shown in non-human primates [49, 50]. The lowest estimates for the core are also consistent with recordings from single neurons in the primary auditory cortex of pre-surgical epileptic patients [2] where about half of excitatory neurons were found to be more finely tuned than in the auditory nerve (i.e., tuning widths < 1 ERB). More extensive electrophysiological measurements in non-human primates however have generally reported wider tuning widths. To explore this further, we digitized cortical tuning widths from seven studies that reported single- or multi-unit tuning width measurements in either core or belt areas in non-human primates (Supplementary Table S3; see Methods section for details). Median neuronal tuning width in non-human primates ranged between 2.8 and 4.9 ERBs in core auditory cortex (depending on the precise assumptions made when converting these tuning width to a format compatible with our measurements, see supplementary methods) and between 3.5 and 6.0 ERBs in belt auditory cortex. While these values are on the low side of our fMRI-fitted tuning width parameter range for the core (0.1 to 14 ERBs) and anterior belt region (3 to 18 ERBs), they are lower than the range obtained for the posterior belt (7 to 23 ERBs). Considering in addition that the non-human primate tuning width estimates may be inflated due to anesthesia [1, but see 51], and that our estimates may reflect sharper tuning in upstream regions, it would thus appear that our fMRI -based tuning width estimates are wider overall than those obtained from direct (invasive) recordings in both humans and non-human primates. This is also the case for adaptation tuning obtained in previous human EEG studies [38, 52, 53]. The EEG-derived tuning width estimates (shown in supplementary Fig. S4), measured from the N1 and/or P2 ERP components, ranged between 15.4 and 20.8 ERBs, with a median of 16.1 ERBs. Considering the 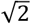bias, this would correspond to a median neuronal tuning width of 11.4 ERBs, which lies in the middle of our fMRI-based tuning width ranges for both the core and belt regions, as would be expected since both regions are thought to contribute to the generation of the N1 and P2 [54]. Therefore, it seems that neuronal frequency selectivity estimates obtained using adaptation with either fMRI or EEG are both wider than those obtained by invasive measurements. This discrepancy may be due to the fact that both BOLD and EEG signals are more sensitive to peri-synaptic activity than to spike rates used in invasive recordings [24, 55]. Intracellular recordings in the auditory system have shown that there is considerable subthreshold synaptic activity in response to frequencies beyond a neuron’s classical frequency receptive field [56]. This subthreshold synaptic activity may contribute to neuronal adaptation and thus lead non-invasive, adaptation-based tuning width estimates to appear wider than invasive extracellular estimates.

Given the large uncertainty of possible neuronal tuning width parameters obtained from our model fits, comparison with invasive recordings remains, of course, highly speculative. The uncertainty in fitted tuning width parameters was linked with the uncertainty in other model parameters, particularly, the tonotopic scatter and the relative strength of neuronal and non-neuronal adaptation, in that, narrower tuning width estimates were associated with greater tonotopic scatter estimates (more variability in the characteristic frequencies at a given cortical location) and stronger non-neuronal non-linearity. Thus, it may be possible to reduce uncertainty in tuning width estimates by fixing some of the other model parameters based on independent information (e.g., independent estimates of tonotopic scatter and/or neuronal adaptation strength) and using a more realistic model of BOLD signal generation [e.g. 57]. For instance, two-photon calcium imaging studies have shown a high degree of fine-scale disorder, in tonotopic organization in mammalian auditory cortex [58, 59], suggesting a relatively high degree of tonotopic scatter, which, according to our model, would imply narrower neuronal tuning width estimates.

## Conclusion

Using fMRI adaptation and computational modelling, we attempted to estimate average single-neuron frequency tuning width in primary and secondary auditory cortex. Our method accounts for several biases due to the macroscopic and non-neural nature of fMRI adaptation. However, there is still uncertainty about whether the measured frequency tuning reflects neuronal tuning within the measured regions, or inherited adaption, despite our efforts to address this question. Further work using more exhaustive stimulus manipulations and more realistic models of cortical adaptation [e.g. 60] may be able to address this issue.

## Methods

This study involved 11 normal-hearing participants with no history of neurological or otological disease (3 males; age range: 29.9 ± 7.4). All participants gave prior-informed written consent. The procedures complied with the Declaration of Helsinki guidelines (Version 6, 2008) and were approved by the Ethics Committee of the University of Nottingham’s School of Medicine. Some of the current data (for conditions not involving adaptation) have been presented previously [7], with the current participants 1 to 11 corresponding to participants 2 to 12 in the previous study (previous participant 1 did not complete all adaptation conditions).

All scanning was performed on a Philips Achieva 7T system with a birdcage transmit, and 32-channel receive coil (Nova Medical, Wilmington, MA). Participants watched a self-chosen silent subtitled movie to stay alert and were told that they could ignore all sound stimuli.

### Functional MRI acquisition, acoustic stimuli and pre-processing

BOLD fMRI data were acquired using a sparse (TR = 7.5 s) gradient-echo EPI sequence (TE = 25 ms, SENSE = 3, FA = 90°) with 1.5 mm isotropic resolution. The acquisition stack consisted of 20 contiguous axial slices oriented parallel to the Sylvian fissure. The experimental stimuli were presented in the silent periods separating successive fMRI acquisitions, starting 1.28 s after the end of the preceding acquisition and continuing for 5 s, until the start of the next acquisition (Fig. 1).

Each 5-s trial contained either no experimental stimuli (“silent” trials), or a train of 10 identical stimuli, presented once every 500 ms (Fig. 1). Each stimulus consisted either of an adaptor-probe pair (AP trials), or the adaptor or probe presented in isolation (A-only or P-only trials). Both the adaptors and probe were narrowband noises with a bandwidth equal to the normal auditory filter bandwidth at the center frequency. The auditory filter bandwidths are based on behavioral notched-noise measurements [3] expressed as equivalent rectangular bandwidths, or ERBs (i.e., the widths of rectangular filters with the same peak gain and total power), and approximated as a linear function, *erb*(*f*) = 24.7 ⋅ (4.37 ⋅ *f* + 1), of the filter center frequencies, *f*. The frequencies of both the adaptors and probes were fixed within each trial. Across trials, the frequency of the probe was fixed at 3.839 kHz, whilst the frequency of the adaptor was varied across seven possible values (0.251, 0.505, 0.899, 1.501, 2.423, 3.839 and 6.009 kHz), spaced evenly on a quasi-logarithmic frequency scale referred to as the ERB-number scale [see 61, and https://en.wikipedia.org/wiki/Equivalent_rectangular_bandwidth], which approximates the human cochlear frequency scale by measuring the number of ERBs below each frequency [62]. For the AP trials, the adaptor frequencies were limited to the central five of these values (0.505, 0.899, 1.501, 2.423, 3.839 kHz), yielding a total of 14 different trial types: silent, P-only, five AP trials and seven A-only trials.

Across different groups of participants, the adaptors were either single-onset 200-ms bursts (Fig. 1B; first five participants, single-onset adaptor group), or short sequences of four 50-ms (Fig. 1C; last six participants, multiple-onset adaptor group). Multiple adaptors were separated by 30ms gaps, which is long enough to elicit separate onset responses [63].. The probe was a single 50-ms burst for all participants. In the AP trials, each probe followed the end of the respective preceding adaptor after a 30-ms silent gap. In the A-only and P-only trials, the adaptor and probe stimuli were presented in the same temporal positions relative to the offset of the preceding fMRI acquisition as in the AP trials. All stimulus bursts were gated on and off with 10-ms quarter-cosine ramps. The stimuli were generated digitally, converted to analogue voltage and presented through MR-compatible insert earphones (Sensimetrics S14) at a root-mean-square level of 70 dB SPL (sound pressure level). They were also equalized to compensate for the frequency transfer characteristics of the earphones, and presented in a continuous background of uniformly exciting noise [filtered to contain equal energy within all cochlear filters; 64] to roughly equate their sensation levels (SLs) across frequencies and participants. The noise was presented at 35 dB SPL per ERB, creating a constant hearing threshold of approximately 29 dB SPL [65].

Data were acquired in 8-min runs consisting of a total of 64 trials each, including four repetitions of each of the 13 non-silent (AP, A and P) trials and 12 repetitions of the silent trials. The SL-equalizing background noise was presented continuously throughout the run duration. The trials were presented in a pseudo-random sequence consisting of four successive random permutations of one of each of the non-silent trials and three silent trials, respectively. Most participants completed a total of six runs, except for participants 2 and 5 who completed eight and four runs, respectively.

The functional images were corrected for distortions due to dynamic B_0_ inhomogeneities [see 7 for details], motion-corrected, high-pass filtered (cut-off 0.01 Hz), converted to percent signal change and finally concatenated across runs. BOLD responses to each of the 13 sound conditions (seven A, five AP and one P condition) were estimated voxelwise by fitting a general linear model to the concatenated functional time series for each subject [see 7 for details].

### Structural MRI acquisition, functional-to-structural alignment and group normalization

A whole-head, high-resolution T1-weighted structural volume was acquired using a phase-sensitive inversion recovery (PSIR) sequence (TR = 15 ms, TE = 6.1 ms, SI = 5000 ms) with 0.6 mm isotropic resolution. This was used both to create a flattened representation of the supratemporal cortical surface around auditory cortex in each subject and hemisphere, and to estimate local intracortical myelin content. An unbiased, semi-quantitative measure of longitudinal relaxation rate (R1), which is indicative of cortical myelination, was obtained by dividing the sign-corrected T1-weighted PSIR volume voxelwise by the corresponding proton-density-weighted volume [see 7]. The resulting R1 volume was segmented to reconstruct individual three-dimensional cortical surfaces at 11 cortical depths [using Freesurfer v5.3, https://surfer.nmr.mgh.harvard.edu/, 66], which, in turn, were used to create the flattened cortical patches (using vistaSoft’s mrVista, http://web.stanford.edu/group/vista/cgi-bin/wiki/index.php/MrVista). To project the functional data onto the cortical patches, the functional volumes were non-linearly registered with the T1-weighed PSIR volume [using an additional T2*-weighted FLASH volume with the same slice prescription as the functional volumes in an intermediate registration step, see 7 for details].

Functional parameter estimates for each of the 13 different sound conditions were also normalized to the MNI305 brain template (fsaverage) using spherical normalization of the 11 cortical surfaces in Freesurfer [67] and mrTools [68]. For all group analyses performed on spherically-normalized data, the left and the right hemispheres of each subject were treated as independent and data were aggregated across the left and the right hemispheres by projecting individual right hemisphere data onto the template’s left hemisphere [see 7 for details].

### *In vivo* parcellation of auditory cortex in individual hemispheres and definition of surface ROIs

To delineate primary from non-primary auditory regions (commonly referred to as “core” and “belt”) in individual hemispheres, we mapped local preferred frequencies, response tuning widths and cortical myelin content onto the individual flat cortical patches [7]. Local preferred frequencies and response tuning widths were derived from the voxelwise frequency response functions based on the BOLD responses to the seven different adaptor-only frequencies (henceforth referred to as “response tuning curves”). Preferred frequencies were obtained by calculating the tuning curve centroids, correcting them for the central bias caused by the finite range of adaptor frequencies, and then averaging them across the central five cortical depths. Response tuning widths were obtained by first recentering the voxelwise tuning curves on their respective centroid frequencies, averaging them across all cortical depths and smoothing them along the cortical surface (2D-Gaussian kernel with 3-mm FWHM, full width at half maximum). The voxelwise, recentred and smoothed tuning curves were then fitted with Gaussian-shaped functions (on the ERB-number scale). Given that the tuning curves were recentered, only the Gaussian scale (determining the size of the peak response) and spread parameters were fitted. Intracortical myelin content was estimated from the PSIR R1 volumes by regressing out local cortical curvature and thickness and averaging across the middle five cortical depths. The preferred frequencies were used to define four contiguous surface regions of interest (ROIs) in each individual hemisphere. Reversals in local tonotopic gradient direction were identified using an unbiased automated procedure [69]. Each individual hemisphere showed a low-frequency reversal along the long axis of Heschl’s gyrus (HG), flanked by two high-frequency reversals anteriorly and posteriorly [see Fig. 2A for an example hemisphere and 7 for all individual hemispheres]. We used these three reversals to define two ROIs corresponding to the two mirror-symmetric tonotopic gradients running across HG (henceforth referred to as “anterior and posterior gradient ROIs”). The anterior gradient ROI substantially overlapped regions of increased frequency selectivity (reduced response tuning width) and increased intracortical myelination observed along the long axis of HG, probably corresponding to the auditory primary, or core, region [see Fig. 2B&C and 7]. The posterior gradient ROI also overlapped the core region, but less so than the anterior gradient ROI. We then defined two additional ROIs posterior and anterior to the two gradient ROIs, representing the secondary, or belt, auditory regions (henceforth referred to as “anterior and posterior belt ROIs”). These were defined as regions significantly responsive to any of the seven adaptor-only frequencies, but not belonging to either of the two gradient ROIs. To show the approximate location of individually-defined ROIs on the group template brain (Fig. 2D), we projected the individual ROI masks to the group template using spherical normalization and computed probability maps across subjects/hemispheres for each ROI. We defined the corresponding group ROIs as the regions where the probability for a given ROI was both larger than for any other ROI and larger than 50%.

### Estimation of ROI-average response and adaptation tuning widths

To derive the ROI-average adaptation tuning widths, we first estimated the adapted response to the probe when it was preceded with different adaptor frequencies (P|A). This was done by taking the difference, AP – A, between the ROI-average responses to each adaptor-probe pair, AP, and adaptor-only stimulus, A, at the corresponding adaptor frequency. We then calculated the amount of adaptation for each adaptor frequency by subtracting the adapted probe responses, P|A, from the unadapted ROI-average response to the probe-only stimulus, P. The resulting differences were expressed as a proportion of the ROI-average probe-only response [(P – P|A)/P]. As only the multiple-adaptor group showed clearly tuned adaptation tuning curves, all further analyses were restricted to this group.

ROI-average response tuning widths were derived similarly to the smoothed local response tuning widths, by first recentering the voxelwise response tuning curves to their bias-corrected centroid frequencies, averaging them across all voxels within a given ROI, and, finally, fitting them with Gaussian-shaped functions (on the ERB-number scale) to estimate the function scale (peak response) and spread.

To measure the widths of the ROI-average adaptation tuning curves (thought to represent the average neuronal tuning width), we fitted them with two types of Gaussian functions. For the first fit, we assumed that adaptation was entirely stimulus-specific (i.e., specific to the probe frequency) and would thus approach zero for sufficiently remote adaptor frequencies. Thus, the fitted functions were simple Gaussians on the ERB-number scale, with free spread and scale parameters, representing tuning width and maximum adaptation amount (when adaptor and probe frequencies matched), respectively. However, in belt ROIs, adaptation seemed to approach a non-zero floor, suggesting that a proportion of the adaptation effect was stimulus-independent. Thus, for the second fit, we fitted Gaussians with a constant offset as an additional free parameter. As ROI-average adaptation was always greatest when the adaptor frequency matched the probe frequency (see Fig. 4B), the location of the function mode was fixed at the probe frequency for both fits. BOLD response tuning widths were obtained from BOLD responses tuning curves using the same methods.

Gaussian functions were fitted both individually to each participant’s hemisphere- and ROI-specific tuning curves (individual fits) and to the ROI tuning curves averaged across all 12 hemispheres in the multiple-adaptor group (group fits). Confidence intervals for the group-average tuning curves (Fig.4) and group-fitted parameters were obtained by bootstrap resampling the 12 hemispheres (with replacement) and fitting each sample average tuning curve in the same way as the group-average curve. Differences in the fitting parameters between ROIs were tested with a linear mixed-effect model (R version 4.0.3, function *lmer* in package *afex*), using the individually fitted parameters after rejection of outliers in the spread parameter using the median absolute deviation method. Post-hoc pairwise comparisons were adjusted for family-wise error rate using Tukey’s range test (R function *lsmeans* in package *emmeans*). All tuning spread parameter estimates are reported as FWHM of the best-fitting Gaussian functions. Goodness of fit and model selection indicators (r^2^, corrected Akaike information criterion, AICc, and Bayes information criterion, BIC) were computed for the group fits with respect to all individual hemisphere tuning curves.

In addition to the ROI-average response and adaptation tuning curves, we computed group-average maps of the adaptor and probe responses (GLM parameter estimates), and of the unnormalized and normalized adaptation effects [(P – P|A) and (P – P|A)/P]. For this, the spherically-normalized individual responses and adaptation effects were first averaged across all cortical depths and then smoothed along the flattened cortical surface with a Gaussian kernel (FWHM=6 mm). The group-average maps were tested for statistical significance against zero using mass univariate one-sample t tests, corrected for multiple tests across all voxels of the left-hemisphere flattened cortical template using an adaptive FDR correction method [70].

### Computational model of fMRI adaptation in tonotopically-organized cortex

To gain a better understanding of how fMRI-based response and adaptation tuning widths are related to the underlying neuronal tuning widths, we compared the measured data with simulations based on a computational model of fMRI adaptation. The model implemented an array of approximately hundred 1.5 mm unidimensional voxels sampling a unidimensional tonotopically-organized cortical strip containing a population of frequency-selective neurons (the exact number of voxels depended on the value of the tonotopic gradient, see below). We used this model to predict the amplitude of the BOLD responses (both at the individual voxel level and averaged across the array) to the adaptor-only (A), probe-only (P) and adaptor-probe (AP) stimuli, (ignoring the temporal dimension of the stimuli and responses for simplicity).

Neurons had Gaussian-shaped, unit-scaled response tuning curves to narrowband stimuli (A and P), *R*_*cell*_ (*f*|*f*_*c*_, *σ*) = *exp*(−(*f*_*c*_ − *f*)^2^/(2*σ*^2^)), where *f* is the stimulus frequency on the ERB-number scale, *f*_*c*_ is the neuron’s characteristic frequency and *σ* determines its tuning width (in ERBs). Numerical values for *σ* are expressed as FWHM throughout the manuscript. *f*_*c*_ was limited to the range of human hearing between 0.02 and 18 kHz. Neuronal adaptation to an adaptor was modelled by decreasing the unadapted response to the probe, *R*_*cell*_ (*P*) = *R*_*cell*_ (*f*_*P*_ |*f*_*c*_, *σ*)multiplicatively by an amount proportional to the adaptor response, *R*_*cell*_ (*A*) = *R*_*cell*_ (*f*_*A*_ |*f*_*c*_, *σ*):

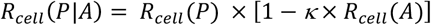

with proportionality constant, *κ*, representing the maximum adaptation strength (i.e. the maximum adaptation that would occur when the adaptor frequency, *f*, matches the neuron’s characteristic frequency, *f*_*c*_, and therefore *R*_*cell*_(*A*) = 1). Both parameters *σ* and *κ* were assumed to be constant across the entire neuron population.

Neuronal characteristic frequencies, *f*_*c*_, were tonotopically organized, increasing, on average, from one end of the cortical strip to the other. To allow for tonotopic scatter, the actual characteristic frequencies (*f*_*c*_) at each point in the cortical strip were allowed to vary randomly around a nominal value, *φ*, which increased linearly in ERB-numbers per millimetre to create a given tonotopic gradient (e.g., 2.5 ERB/mm). Thus, at any point along the strip, *f*_*c*_ was allowed to vary according to a truncated Gaussian distribution centred on the local value of *φ*:

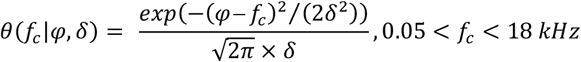

The spread parameter, *δ*, of the scatter distribution controls the degree of randomness in local characteristic frequency values, with zero scatter (*δ* = 0) corresponding to a perfectly ordered spatial progression of characteristic frequencies, and infinite scatter (*δ* = ∞) corresponding to a completely random distribution of characteristic frequencies across the strip. Numerical values for *δ* are expressed as FWHM in ERBs throughout the manuscript. Note that changing the amount of scatter also changed the slope of the average characteristic frequency progression, or tonotopic gradient, across the strip.

The neuron population was sampled using an array of equally-spaced, but overlapping, 1.5 mm voxels. Although voxels overlapped, they were spaced in such a way that there was an integer number of contiguous voxels spanning a range of nominal characteristic frequencies (*φ*) between about 0.05 and 16 kHz. The exact range was adjusted as required to create a given tonotopic gradient (e.g., 2.5 ERB/mm).

The BOLD response to a given stimulus (A, P or AP) in a given voxel was obtained by, first, integrating the neuronal responses to the stimulus across the full distribution of characteristic frequencies, *f*_*c*_, of neurons contained within the voxel, then convolving with a hemodynamic point spread function (PSF) and, finally, applying a compressive function to account for non-neuronal (neurovascular) non-linearities [e.g., 28].

The average neuronal response in a voxel (i.e. before applying the non-linearity and convolving with the PSF), *R*_*voxel*_ (*f*), to an A or P stimulus centered at frequency *f* depended on the distribution of characteristic frequencies within the voxel, and therefore on both the value of the spatial gradient of *φ* and on the scatter parameter *δ* :

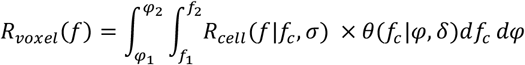

where *φ*_*l*_ and *φ*_2_ are the lower and upper bounds of the nominal characteristic frequencies within the voxel, as determined by the tonotopic gradient, and *f*_*l*_ and *f*_2_ are the bounds of the hearing range (0.01-18 kHz), all expressed in ERBs.

A voxel’s average adapted neuronal response to the probe (i.e., following an adaptor stimulus) was similarly computed by integrating the adapted neuronal responses *R*_*cell*_(*P*|*A*) within the voxel:

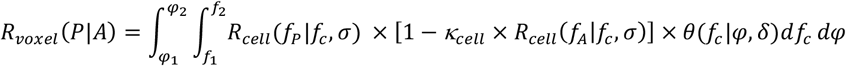

Finally, the voxel’s average neuronal response to an adaptor-probe pair, *R*_*voxel*_(*AP*), was computed as the sum of *R*_*voxel*_(*A*) and *R*_*voxel*_(*P*|*A*).

The hemodynamic PSF was modelled by a spatial Gaussian convolution kernel, *g*, with an FWHM of 2mm. Thus, a voxel’s BOLD response to a given stimulus, S, was given by *R*_*BOLD*_ (*S*) = *g* ∘ *s*(*R*_*voxel*_ (*S*)) To avoid edge effects when convolving responses with the hemodynamic PSF (see below), we added mirror images of the cortical strip and voxel array on each side of it (i.e. with characteristic frequency progressions opposite to that in the main array, resulting in frequency reversals). Responses from voxels in these duplicated arrays were otherwise ignored.

Neurovascular non-linearity was modelled as a compressive function, *s*, applied to the neuronal voxel response of each type of stimulus (A, P or AP) and implemented as the positive half of a re-centered logistic function, such that *S*(0) = 0, and scaled to have unit derivative at 0 and to reach a saturated BOLD level γ:

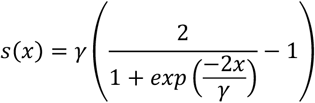

In actual experimental conditions, the BOLD response to the adapted probe (corresponding to *R*′′_*voxel*_ (*P*|*A*) in the model), cannot be observed on its own, because, due to the temporal sluggishness of the hemodynamic response, it overlaps with the response to the preceding adaptor (corresponding to *R*_*BOLD*_ (*A*)). For comparison with the experimental fMRI data, we thus estimated *R*_*BOLD*_ (*P*|*A*) by calculating the difference between the responses to the respective adaptor probe pair and adaptor-alone stimuli, 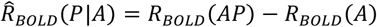. The associated adaptation effect was then obtained by calculating the difference between the unadapted and adapted probe responses, 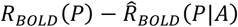, as in the experiment.

The modelled voxelwise adaptor-only BOLD responses and fMRI adaptation effects at different adaptor frequencies were then used to construct array-average BOLD responses and BOLD adaption tuning curves and derive the associated tuning widths. This was done in exactly the same way as for the corresponding experimental data. In particular, to create array-average response tuning curves (based on the adaptor-only responses), the modelled voxelwise response tuning curves were first recentered on their estimated bias-corrected centroids. Tuning widths were then derived by fitting the tuning curves with Gaussian functions and the adaptation tuning curves were fitted twice –with or without an added constant offset to represent any stimulus-independent adaptation.

### Exploring and fitting the model

We first explored the model’s behavior informally by manually setting its four free parameters (neuronal tuning spread, *σ*, tonotopic scatter, *δ*, neuronal adaptation strength, *κ*, BOLD saturation point, γ) to specific values (see Fig. 6). The nominal neuronal characteristic frequency gradient was set to 2.5 ERBs/mm, close to the tonotopic gradient measured in our data (see below). To obtain smooth-looking tuning curves, we simulated responses for 50 adaptor frequencies ranging between 0.19 and 7.83 kHz, rather than just the seven frequencies used in the experiment.

Then, we fitted the model parameters by minimizing the sum of squared differences between the measured group-average and ROI-average response and adaptation tuning curves and the corresponding array-average model tuning curves (with each ROI modelled as a separate tonotopic array), simulated for the seven adaptor frequencies and five adaptor-probe pairs used in the experiment. Response and adaptation tuning curves from the two gradient and the two belt ROIs were fitted jointly using the constrained non-linear programming solver *fmincon* in Matlab (with the default interior-point algorithm). During fitting, both the modelled and measured tuning curves were normalized to the peak of the response tuning curve. This was to avoid having to include a scaling parameter to match the (arbitrary) units of the modelled responses to those of the measured ones. The response tuning curves contained a maximum of 17 data points (due to recentering voxel tuning curves according to their bias-corrected centroid) and the adaptation tuning curves contained five points, yielding a total of 22 data points for each ROI. To equate the contribution of each type of tuning curve (response versus adaptation) to the fit, the cost function (sum of squared errors) was weighted inversely to the tuning curve area (given by the sum of all its data points). In each fit, the neuronal tuning spread, *σ*, the tonotopic scatter, *δ*, and the neuronal adaptation strength, *κ*, were allowed to vary between ROIs, whilst the BOLD saturation level, γ, was fixed across ROIs, resulting in a total of 10 (3 ×3 +1) free parameters for each fit. *σ, δ* and γ were constrained to be positive and *К* was constrained to range between 0 and 1. The model’s nominal tonotopic gradient strength was set to the gradient strength observed in the actual data in each ROI, averaged across the 22 hemispheres. The gradient strength was computed in each ROI of each individual hemisphere by averaging the local preferred-frequency (bias-corrected centroid, see above) gradient strength measured at each face of the 3D cortical mesh [71] contained in each ROI.

The model was fitted 1000 times in order to obtain the distributions and joint distributions of best-fitting parameter values. For each fit, the initial parameter values were drawn randomly from predefined intervals ([0 50] ERBs for *σ* and *δ*, [0 1] for *К*, and κ 1 +∞] for γ). The interval for γ corresponded to a BOLD saturation point between 1.79 and +∞ % signal change when taking into account the tuning curve normalization constant averaged across ROIs).

### Comparisons with tuning width estimates from the human and non-human primate electrophysiological literature

To compare our fMRI-adaptation-based frequency tuning width estimates with neuronal tuning widths measured electrophysiologically, we digitized data from seven studies that reported single- or multi-unit excitatory frequency tuning width from auditory neurons in core and/or belt fields using different non-human primate species in either anesthesized or unanesthesized recording conditions [see supplementary Table 1; 1, 72-77], and from 3 EEG studies that used a frequency adaptation paradigm similar to ours and measured adaptation tuning of the N1 and/or P2 ERP components [52, 53, 78].

To make the non-human primate measurements as comparable to our fMRI-based tuning width estimates as possible, we converted the digitized primate tuning widths, originally given either as iso-response quality factors or as iso-response tuning widths in octaves, to iso-intensity FWHMs based on an assumed Gaussian tuning shape on the human ERB-number scale (see supplementary methods section for details).

Digitized frequency adaptation data from the three EEG studies were re-expressed as relative decrease in the unadapted N1 response amplitude (or N1/P2 amplitude difference) to a probe presented alone, when this probe was preceded by adaptors at different frequencies (normalized adaptation in % as in our Fig. 3F and 4B), and the adaptor-probe frequency difference was re-expressed in cochlear ERBs. The resulting adaptation tuning curves (shown in supplementary Fig. S4) were fitted with simple Gaussian functions centered on the probe frequency to obtain the tuning FWHM.

## References

1. Bartlett EL, Sadagopan S, Wang X. Fine frequency tuning in monkey auditory cortex and thalamus. J Neurophysiol. 2011;106(2):849–59.

2. Bitterman Y, Mukamel R, Malach R, Fried I, Nelken I. Ultra-fine frequency tuning revealed in single neurons of human auditory cortex. Nature. 2008;451(7175):197–201.

3. Glasberg BR, Moore BC. Derivation of auditory filter shapes from notched-noise data. Hear Res. 1990;47(1-2):103–38.

4. Patterson RD, Nimmo-Smith I. Off-frequency listening and auditory-filter asymmetry. J Acoust Soc Am. 1980;67(1):229–45.

5. Rosen S, Baker RJ. Characterising auditory filter nonlinearity. Hear Res. 1994;73(2):231–43.

6. Shera CA, Guinan JJ, Jr., Oxenham AJ. Revised estimates of human cochlear tuning from otoacoustic and behavioral measurements. Proc Natl Acad Sci U S A. 2002;99(5):3318–23.

7. Besle J, Mougin O, Sanchez-Panchuelo RM, Lanting C, Gowland P, Bowtell R, et al. Is human auditory cortex organization compatible with the monkey model? Contrary evidence from ultra-high-field functional and structural MRI. Cereb cortex. 2019;29(1):410–28.

8. Thomas JM, Huber E, Stecker GC, Boynton GM, Saenz M, Fine I. Population receptive field estimates of human auditory cortex. Neuroimage. 2015;105:428–39.

9. De Martino F, Moerel M, Xu J, van de Moortele PF, Ugurbil K, Goebel R, et al. High-Resolution Mapping of Myeloarchitecture In Vivo: Localization of Auditory Areas in the Human Brain. Cereb cortex. 2015;25(10):3394–405.

10. Moerel M, De Martino F, Formisano E. Processing of natural sounds in human auditory cortex: tonotopy, spectral tuning, and relation to voice sensitivity. J Neurosci. 2012;32(41):14205–16.

11. Collins CE, Turner EC, Sawyer EK, Reed JL, Young NA, Flaherty DK, et al. Cortical cell and neuron density estimates in one chimpanzee hemisphere. Proc Natl Acad Sci U S A. 2016;113(3):740–5.

12. Collins CE, Airey DC, Young NA, Leitch DB, Kaas JH. Neuron densities vary across and within cortical areas in primates. Proc Natl Acad Sci U S A. 2010;107(36):15927–32.

13. Kamitani Y, Tong F. Decoding the visual and subjective contents of the human brain. Nat Neurosci. 2005;8(5):679–85.

14. Barron HC, Garvert MM, Behrens TE. Repetition suppression: a means to index neural representations using BOLD? Philos Trans R Soc Lond B Biol Sci. 2016;371(1705).

15. Grill-Spector K, Henson R, Martin A. Repetition and the brain: neural models of stimulus-specific effects. Trends Cogn Sci. 2006;10(1):14–23.

16. Huettel SA, Obembe OO, Song AW, Woldorff MG. The BOLD fMRI refractory effect is specific to stimulus attributes: evidence from a visual motion paradigm. Neuroimage. 2004;23(1):402–8.

17. Nishida S, Sasaki Y, Murakami I, Watanabe T, Tootell RB. Neuroimaging of direction-selective mechanisms for second-order motion. J Neurophysiol. 2003;90(5):3242–54.

18. Weigelt S, Limbach K, Singer W, Kohler A. Orientation-selective functional magnetic resonance imaging adaptation in primary visual cortex revisited. Hum Brain Mapp. 2012;33(3):707–14.

19. Tootell RB, Hadjikhani NK, Vanduffel W, Liu AK, Mendola JD, Sereno MI, et al. Functional analysis of primary visual cortex (V1) in humans. Proc Natl Acad Sci U S A. 1998;95(3):811–7.

20. Fang F, Murray SO, Kersten D, He S. Orientation-tuned FMRI adaptation in human visual cortex. J Neurophysiol. 2005;94(6):4188–95.

21. Lee HA, Lee SH. Hierarchy of direction-tuned motion adaptation in human visual cortex. J Neurophysiol. 2012;107(8):2163–84.

22. Krekelberg B, Boynton GM, van Wezel RJ. Adaptation: from single cells to BOLD signals. Trends Neurosci. 2006;29(5):250–6.

23. Larsson J, Solomon SG, Kohn A. fMRI adaptation revisited. Cortex. 2016;80:154–60.

24. Bartels A, Logothetis NK, Moutoussis K. fMRI and its interpretations: an illustration on directional selectivity in area V5/MT. Trends Neurosci. 2008;31(9):444–53.

25. Verhoef BE, Kayaert G, Franko E, Vangeneugden J, Vogels R. Stimulus similarity-contingent neural adaptation can be time and cortical area dependent. J Neurosci. 2008;28(42):10631–40.

26. Scholes C, Palmer AR, Sumner CJ. Forward suppression in the auditory cortex is frequency-specific. Eur J Neurosci. 2011;33(7):1240–51.

27. Vogels R. Sources of adaptation of inferior temporal cortical responses. Cortex. 2016;80:185–95.

28. Magri C, Logothetis NK, Panzeri S. Investigating static nonlinearities in neurovascular coupling. Magn Reson Imaging. 2011;29(10):1358–64.

29. Malach R. Targeting the functional properties of cortical neurons using fMR-adaptation. Neuroimage. 2012;62(2):1163–9.

30. Carandini M, Ferster D. A tonic hyperpolarization underlying contrast adaptation in cat visual cortex. Science. 1997;276(5314):949–52.

31. Brosch M, Scheich H. Tone-sequence analysis in the auditory cortex of awake macaque monkeys. Exp Brain Res. 2008;184(3):349–61.

32. Sawamura H, Orban GA, Vogels R. Selectivity of neuronal adaptation does not match response selectivity: a single-cell study of the FMRI adaptation paradigm. Neuron. 2006;49(2):307–18.

33. Liu Y, Murray SO, Jagadeesh B. Time course and stimulus dependence of repetition-induced response suppression in inferotemporal cortex. J Neurophysiol. 2009;101(1):418–36.

34. De Baene W, Vogels R. Effects of adaptation on the stimulus selectivity of macaque inferior temporal spiking activity and local field potentials. Cereb cortex. 2010;20(9):2145–65.

35. Abbott LF, Varela JA, Sen K, Nelson SB. Synaptic depression and cortical gain control. Science. 1997;275(5297):220–4.

36. Wehr M, Zador AM. Synaptic mechanisms of forward suppression in rat auditory cortex. Neuron. 2005;47(3):437–45.

37. Reyes AD. Synaptic short-term plasticity in auditory cortical circuits. Hear Res. 2011;279(1-2):60–6.

38. Briley PM, Krumbholz K. The specificity of stimulus-specific adaptation in human auditory cortex increases with repeated exposure to the adapting stimulus. J Neurophysiol. 2013;110(12):2679–88.

39. Duque D, Wang X, Nieto-Diego J, Krumbholz K, Malmierca MS. Neurons in the inferior colliculus of the rat show stimulus-specific adaptation for frequency, but not for intensity. Sci Rep. 2016;6:24114.

40. Mill R, Coath M, Wennekers T, Denham SL. A neurocomputational model of stimulus-specific adaptation to oddball and Markov sequences. PLoS Comput Biol. 2011;7(8):e1002117.

41. Inan S, Mitchell T, Song A, Bizzell J, Belger A. Hemodynamic correlates of stimulus repetition in the visual and auditory cortices: an fMRI study. Neuroimage. 2004;21(3):886–93.

42. Heckman GM, Bouvier SE, Carr VA, Harley EM, Cardinal KS, Engel SA. Nonlinearities in rapid event-related fMRI explained by stimulus scaling. Neuroimage. 2007;34(2):651–60.

43. Zhou J, Benson NC, Kay KN, Winawer J. Compressive Temporal Summation in Human Visual Cortex. J Neurosci. 2018;38(3):691–709.

44. Zhang N, Zhu XH, Chen W. Investigating the source of BOLD nonlinearity in human visual cortex in response to paired visual stimuli. Neuroimage. 2008;43(2):204–12.

45. Bao PL, Purington CJ, Tjan BS. Using an achiasmic human visual system to quantify the relationship between the fMRI BOLD signal and neural response. Elife. 2016;5.

46. Evans EF. The frequency response and other properties of single fibres in the guinea-pig cochlear nerve. J Physiol. 1972;226(1):263–87.

47. Keliris GA, Li Q, Papanikolaou A, Logothetis NK, Smirnakis SM. Estimating average single-neuron visual receptive field sizes by fMRI. Proc Natl Acad Sci U S A. 2019;116(13):6425–34.

48. Zulfiqar I, Havlicek M, Moerel M, Formisano E. Predicting neuronal response properties from hemodynamic responses in the auditory cortex. Neuroimage. 2021;244:118575.

49. Kosaki H, Hashikawa T, He J, Jones EG. Tonotopic organization of auditory cortical fields delineated by parvalbumin immunoreactivity in macaque monkeys. J Comp Neurol. 1997;386(2):304–16.

50. Morel A, Garraghty PE, Kaas JH. Tonotopic organization, architectonic fields, and connections of auditory cortex in macaque monkeys. J Comp Neurol. 1993;335(3):437–59.

51. Moshitch D, Las L, Ulanovsky N, Bar-Yosef O, Nelken I. Responses of neurons in primary auditory cortex (A1) to pure tones in the halothane-anesthetized cat. J Neurophysiol. 2006;95(6):3756–69.

52. Picton TW, Woods DL, Proulx GB. Human auditory sustained potentials. II. Stimulus relationships. Electroencephalogr Clin Neurophysiol. 1978;45(2):198–210.

53. Butler RA. Effect of changes in stimulus frequency and intensity on habituation of the human vertex potential. J Acoust Soc Am. 1968;44(4):945–50.

54. Yvert B, Fischer C, Bertrand O, Pernier J. Localization of human supratemporal auditory areas from intracerebral auditory evoked potentials using distributed source models. Neuroimage. 2005;28(1):140–53.

55. Goense JB, Logothetis NK. Neurophysiology of the BOLD fMRI signal in awake monkeys. Curr Biol. 2008;18(9):631–40.

56. Xie R, Gittelman JX, Pollak GD. Rethinking tuning: in vivo whole-cell recordings of the inferior colliculus in awake bats. J Neurosci. 2007;27(35):9469–81.

57. Buxton RB. Dynamic models of BOLD contrast. Neuroimage. 2012;62(2):953–61.

58. Rothschild G, Nelken I, Mizrahi A. Functional organization and population dynamics in the mouse primary auditory cortex. Nat Neurosci. 2010;13(3):353–60.

59. Bandyopadhyay S, Shamma SA, Kanold PO. Dichotomy of functional organization in the mouse auditory cortex. Nat Neurosci. 2010;13(3):361–8.

60. Kudela P, Boatman-Reich D, Beeman D, Anderson WS. Modeling Neural Adaptation in Auditory Cortex. Front Neural Circuits. 2018;12:72.

61. Moore BC, Glasberg BR. Suggested formulae for calculating auditory-filter bandwidths and excitation patterns. J Acoust Soc Am. 1983;74(3):750–3.

62. Greenwood DD. A cochlear frequency-position function for several species--29 years later. J Acoust Soc Am. 1990;87(6):2592–605.

63. Gutschalk A. MEG Auditory Research. In: Supek S, Aine CJ, editors. Magnetoencephalography: From Signals to Dynamic Cortical Networks. Cham: Springer International Publishing; 2019. p. 1–35.

64. Glasberg BR, Moore BC. Frequency selectivity as a function of level and frequency measured with uniformly exciting notched noise. J Acoust Soc Am. 2000;108(5 Pt 1):2318–28.

65. Hawkins JE, Stevens SS. The Masking of Pure Tones and of Speech by White Noise. The Journal of the Acoustical Society of America. 1950;22(1):6–13.

66. Dale AM, Fischl B, Sereno MI. Cortical surface-based analysis. I. Segmentation and surface reconstruction. Neuroimage. 1999;9(2):179–94.

67. Fischl B, Sereno MI, Tootell RB, Dale AM. High-resolution intersubject averaging and a coordinate system for the cortical surface. Hum Brain Mapp. 1999;8(4):272–84.

68. Gardner JL, Merriam EP, Schluppeck D, Besle J, Heeger DJ. mrTools: Analysis and visualization package for functional magnetic resonance imaging data.. 2018.

69. Schönwiesner M, Dechent P, Voit D, Petkov CI, Krumbholz K. Parcellation of Human and Monkey Core Auditory Cortex with fMRI Pattern Classification and Objective Detection of Tonotopic Gradient Reversals. Cereb cortex. 2015;25(10):3278–89

70. Benjamini Y, Krieger AM, Yekutieli D. Adaptive linear step-up procedures that control the false discovery rate. Biometrika. 2006;93(3):491–507.

71. Mancinelli C, Livesu M, Puppo E. A comparison of methods for gradient field estimation on simplicial meshes. Computers & Graphics-Uk. 2019;80:37–50.

72. Cheung SW, Bedenbaugh PH, Nagarajan SS, Schreiner CE. Functional organization of squirrel monkey primary auditory cortex: responses to pure tones. J Neurophysiol. 2001;85(4):1732–49.

73. Kajikawa Y, de La Mothe L, Blumell S, Hackett TA. A comparison of neuron response properties in areas A1 and CM of the marmoset monkey auditory cortex: tones and broadband noise. J Neurophysiol. 2005;93(1):22–34.

74. Pelleg-Toiba R, Wollberg Z. Tuning properties of auditory cortex cells in the awake squirrel monkey. Exp Brain Res. 1989;74(2):353–64.

75. Philibert B, Beitel RE, Nagarajan SS, Bonham BH, Schreiner CE, Cheung SW. Functional organization and hemispheric comparison of primary auditory cortex in the common marmoset (Callithrix jacchus). J Comp Neurol. 2005;487(4):391–406.

76. Recanzone GH, Guard DC, Phan ML. Frequency and intensity response properties of single neurons in the auditory cortex of the behaving macaque monkey. J Neurophysiol. 2000;83(4):2315–31.

77. Recanzone GH, Schreiner CE, Merzenich MM. Plasticity in the frequency representation of primary auditory cortex following discrimination training in adult owl monkeys. J Neurosci. 1993;13(1):87–103.

78. Briley PM, Breakey C, Krumbholz K. Evidence for pitch chroma mapping in human auditory cortex. Cereb cortex. 2013;23(11):2601–10.

