## supplementary for "Can single-neuron frequency tuning in human auditory cortex be quantified through fMRI adaptation?"

### Supplementary methods

#### A. Searchlight fit of Gaussian models to voxelwise FMRI adaptation and BOLD response tuning curves

To estimate voxelwise FMRI adaptation and BOLD response tuning widths, we fitted both simple Gaussians and Gaussians with offset to each voxel's group-average adaptation and response tuning curves (averaged across left and right hemispheres for all participants, irrespective of the type of adaptor, single-onset or multiple-onset). For fitting, the tuning curves were averaged across all cortical depths and smoothed along the cortical surface (FWHM = 6 mm). Before smoothing, we recentered each voxel's response tuning curve to its preferred frequency, estimated by the bias-corrected tuning-curve centroid.

#### B. Conversion of single/multi-unit tuning widths from electrophysiological studies

We digitized 19 cortical tuning width datasets across seven published studies, obtained in various areas of non-human primate auditory cortex using single- or multi-unit recordings (see Table S3 for a summary). Five studies (Cheung et al. 2001; Pelleg-Toiba and Wollberg 1989; Philibert et al. 2005; Recanzone et al. 2000; Recanzone et al. 1993) reported scatter plots of characteristic frequencies and corresponding iso-response (IR) tuning widths for the entire population of recorded units (or multi-units). An IR tuning curve measures the stimulus intensity required to elicit a minimum response level (above spontaneous spiking rate) as a function of stimulus frequency (see section B.2 for a more formal definition). The width of this iso-response tuning curve is then measured at a certain intensity level, usually 10 or 40 dB above the iso-response tuning curve minimum (i.e. at the characteristic frequency). Across these five studies, there were 13 datasets recorded from four different cortical areas (A1, R, CM or L), reporting IR tuning widths at either (or both) of two intensity levels (10 or 40 dB) and expressed either in octaves or as quality, or Q, factors (i.e., the ratio of the characteristic frequency over the tuning width, both in linear units, e.g. kHz). First, we converted all IR tuning widths to human cochlear ERB-number units (see section B.1 below for details), and then, we converted them to iso-intensity (II) units. Contrary to an IR tuning curve, an II tuning curve, such as the BOLD response and adaptation tuning curves measured in the current study, measures the response size as a function of the stimulus frequency elicited by a constant stimulus intensity across frequencies. Thus, converting the electrophysiological IR tuning widths to II units is necessary to make them comparable to our fMRI-based tuning width estimates (see sections B.2 and B.3 below for details). Finally, we averaged tuning widths across neuronal units for each dataset (Table S3). A sixth study (Kajikawa et al. 2005) reported IR tuning width histograms without the corresponding characteristic frequencies (four datasets, at 10 and 40 dB intensity levels, recorded from areas A1 and CM). For this study we first converted the histogram bin edges (given in octaves) to II tuning widths using the same methods as for the other five studies (except that the characteristic frequencies were unknown, and so, we used our study's probe frequency, 3.839 kHz instead) and then calculated the histogram average for each dataset. The seventh and most recent study (Bartlett et al. 2011), reported II tuning widths, expressed as equivalent rectangular bandwidths and averaged across neurons (two datasets: one with stimuli presented 10 dB above firing threshold and the other with stimuli presented at the intensity giving the largest response). For both datasets, II tuning widths were given separately for sharply or broadly tuned neurons (tuning widths below or above 0.2 oct, respectively). We combined sharp and broad tuning widths through a weighted average, using the respective population numbers given in the study. Average equivalent rectangular bandwidths for the two datasets were converted to FWHM by multiplication with $2\sqrt{2 \times ln2}/\sqrt{2\pi} = 0.9394$ (based on the assumption that the respective tuning curves were Gaussian).

Averaging across the 19 datasets was performed separately for the core and belt regions (Table S3). As the average tuning width estimates varied widely across studies, and estimates for the belt region were based on only four datasets across two studies (Kajikawa et al. 2005; Recanzone et al. 2000), we instead calculated the belt average tuning width by taking the ratio of the core and belt tuning widths in these two studies, and then multiplying this ratio with the core tuning width averaged across all seven studies.

#### B.1 Conversion of octave tuning widths to human ERB-number units

The ERB number scale (here referred to as NERB scale) measures the number of equivalent rectangular auditory filter bandwidths, or ERBs, located below a given frequency, $f$, in kHz (Glasberg and Moore 1990), and is given by

$$nerb(f) = 21.4 \cdot \log_{10}(4.37 \cdot f + 1) \qquad \text{(Eq. 1).}$$

Using the inverse of this function, a linear tuning width (in kHz), $\Delta f$, can be rewritten as $\left(10^{nerb(f_c)+\Delta nerb/2} - 10^{nerb(f_c)-\Delta nerb/2}\right)/4.37$, where $f_c$ is the characteristic frequency, and $\Delta nerb$ is the tuning width in NERB units. Solving this for $\Delta nerb$ yields

$$\Delta nerb = 42.72 \cdot \left( \log_{10} \left( \frac{4.37 \cdot \Delta f}{4.37 \cdot f_c + 1} + \sqrt{\left( \left( \frac{4.37 \cdot \Delta f}{4.37 \cdot f_c + 1} \right)^2 + 4 \right)} \right) - log_{10}(2) \right) \qquad \text{(Eq. 2).}$$

The linear tuning widths ($\Delta f$) were converted from tuning widths originally given either in octaves or as quality factors, depending on the dataset. In either case, the conversion is straightforward when $f_c$ is known.

Figure S3 shows all digitized data points from the 13 datasets across the five studies that measured IR tuning widths as a function of characteristic frequency, converted to NERB units. All datasets except two (from belt area L) displayed a negative relationship between tuning width and characteristic frequency above 0.5 kHz. Since we were interested in the neuronal tuning width at the current probe frequency (3.839 kHz), only the tuning widths of neurons with characteristic frequencies within one octave of this frequency (see dashed lines in Fig. S2A) were considered further.

#### B.2 Conversion of iso-response to iso-intensity tuning widths

In all but one of the seven primate studies (Bartlett et al. 2011), the reported tuning widths were derived from "iso-response" (IR) tuning curves, which measure the stimulus intensity $I(f)$ required to elicit a fixed response criterion, $R_0$, as a function of the stimulus frequency, $f$. The width of IR tuning curves is measured at a given intensity level, $L_I$, above the minimum intensity, $I_{min} = I(f_c)$, measured at the preferred, or "characteristic", frequency, $f_c$. Across all six digitized studies, the IR tuning widths were measured at either $L_I$ = 10 or $L_I$ = 40 dB above $I_{min}$. In contrast, both our fMRI-based tuning widths and the tuning widths reported by Bartlett et al. (2011) were derived from "iso-intensity" (II) tuning curves, which measure the response level as a function of stimulus frequency, $R(f)$, elicited at a fixed stimulus intensity $I_0$ (70 dB SPL in our study). The width of II tuning curves can be measured either at a given response amplitude below the peak response, $R_{peak} = R(f_c)$ (e.g., at the half-maximum response, as in our study, i.e. $R_{peak}/2$), or, as in Bartlett et al. (2011), as equivalent rectangular bandwidth, that is, the total tuning curve area divided by $R_{peak}$.

The relationship between IR and II tuning widths is not trivial and depends on the shape of the input/output (IO) function relating the measured response amplitude, $R$ (neuronal or BOLD), to the stimulus intensity, $I$. When the IO function is linear, that is $R$ is proportional to $I$, IR and II tuning curves are inversely related through a common frequency transfer function, $W(f)$, with $R(f)$ proportional to $W(f)$ and $I(f)$ proportional to $1/W(f)$ (see section B.3 below). As a result, the associated tuning widths are equivalent as long as they are measured at corresponding intensity and response levels, respectively. For instance, II tuning widths measured at the half-maximum response, $R_{peak}/2$, would be the same as IR tuning widths measured at twice the minimum intensity, $2 \cdot I_{min}$ (corresponding to $L_I$ = 3 dB above $I_{min}$). Thus, to make the neuronal and fMRI-based tuning widths comparable, we would merely need to correct for the difference between the intensity level, $L_I$, at which the neuronal IR tuning widths were measured (10 or 40 dB) and the response level $L_R$ at which we measured the BOLD tuning width (half-maximum and thus $L_R$ = 3dB below $R_{peak}$). Under the assumption that the common underlying transfer function $W(f)$ is Gaussian-shaped on the NERB scale, this correction can be accomplished by multiplying the neuronal (IR) tuning widths by a factor $\sqrt{\ln(2)/\ln(10^{L_I/10})}$, where $L_I$ is either 10 or 40 dB, depending on the dataset (see section B.3).

When the IO function is non-linear, IR and II tuning width estimates taken at equivalent response and intensity levels will differ (Eustaquio-Martin and Lopez-Poveda 2011; Lopez-Poveda and Eustaquio-Martin 2013). However, if the IO function shape is known, this can be corrected for. Here, we compare different corrections based on the assumption that the IO function can be described as a compressive power law. In Section B.3 (below), we show that, under the further assumption that the BOLD response is linearly related to the underlying neuronal response (Wan et al. 2006), and thus BOLD and neuronal responses are subject to the same power-law non-linearity, with the same compression exponent, $\alpha$, the primate IR tuning widths must be multiplied by a modified factor $\sqrt{\ln(2)/(\alpha \cdot \ln(10^{L_I/10}))}$ compared to the linear case (see section B.3). We set the power law exponent, $\alpha$, to either 0.3 or 0.1. These values characterize the approximate shape of the relationship between the perceptual intensity, or "loudness", of sounds and their physical intensity, often referred to as the "loudness function". On average across all intensities, the loudness function has been shown to exhibit a compressive power-law exponent of about 0.3, increasing to around 1 towards threshold (i.e., an approximately linear relationship; Hellman and Zwislocki 1961) and decreasing to around 0.1 at middle and higher intensities (Buus and Florentine 2001). The idea that the non-linearity of auditory cortical responses may be characterized by similar power-law exponents as the loudness function is supported by findings from several previous fMRI studies suggesting that the BOLD response size in auditory cortex is more closely related to loudness than to physical sound intensity (review in Uppenkamp and Rohl 2014).

##### B.3 Relation between iso-response and iso-intensity tuning widths

For a linear filter (i.e. when the IO function is linear), the filter response, $R(f,I)$, as a function of the stimulus frequency, $f$, and intensity, $I$, is given by the product of the filter's frequency transfer function, $W(f)$, and $I$: $R(f,I) = W(f) \cdot I$. As a result, the filter's IR and II tuning characteristics are related through a simple inverse relationship, with $I(f) = R_0/W(f)$ (IR) and $R(f) = W(f) \cdot I_0$ (II), where $R_0$ is the fixed response criterion used to measure the IR curve (usually corresponding to the smallest detectable response above spontaneous firing), and $I_0$ is the fixed stimulus intensity used to measure the II curve. As a result, IR and II tuning widths will be equal as long as they are measured at the same intensity and response levels, respectively, relative to the minimum intensity, $I_{min}$, and maximum response amplitude, $R_{peak}$ (both assumed to be located at the characteristic frequency, $f_c$). Specifically, if the II tuning width is measured at a response amplitude of $R_{peak} \cdot \theta$, with $\theta < 1$ (e.g., $\theta = 0.5$ for the half-maximum tuning width), corresponding to a response level $L_R = 10 \cdot log10(\theta)$ ($L_R$ = 3 dB below $R_{peak}$ in this example), then the equivalent IR tuning width would be measured at an intensity of $1/\theta \cdot I_{min}$, or an intensity level $L_I = 10 \cdot log10(1/\theta)$ ($L_I$ = 3 dB above $I_{min}$ in this example). Since the digitized IR widths were in fact reported at $L_I$ = 10 or 40 dB, we need to find the relationship between IR widths measured at $L_I$ and II widths measured at $L_R$ = 3 dB (corresponding to the FWHM used to characterize our fMRI-based II tuning widths). Under the assumption that $W(f)$ is Gaussian-shaped, $W(f) = exp(-(f - f_c)^2/2\sigma^2)$, the IR tuning width at an intensity level $L_I$ above $I_{min}$ (or equivalently, the II tuning width at a response level $L_R$ below $R_{peak}$) is given by $\Delta f(L) = 2\sigma\sqrt{2ln\left(10^{L_{I/R}/10}\right)}$. Therefore, the ratio between II tuning widths measured at 3 dB and IR tuning widths measured at $L_I$ = 10 or 40 dB is $2\sigma\sqrt{2ln(10^{3/10})}/2\sigma\sqrt{2ln(10^{L_I/10})} \approx \sqrt{\ln(2)/\ln(10^{L_I/10})}$.

When the IO function is non-linear, the relationship between IR and II tuning characteristics is more complex. Even assuming that the neuronal and BOLD responses are affected by the same non-linearity (i.e., the BOLD response is linear with respect to the neuronal response; see Section B.2), the IR curves from neuronal recordings and II curves from BOLD recordings would be differently affected by the non-linearity of the IO function. Under the simplifying assumption that the non-linearity is characterized by a compressive power law with exponent $\alpha < 1$ (see Section B.2), and that it follows (rather than precedes) frequency filtering, the non-linear intensity-response relationship becomes $R(f,I) = (W(f) \cdot I)^{\alpha}$. In this case, the IR curve still reflects the original (Gaussian) transfer function $W(f)$ because $I(f) = \sqrt[\alpha]{R_0}/W(f)$ and $\sqrt[\alpha]{R_0}$ is a constant. In contrast, the II curve is compressed, $R(f) = W(f)^{\alpha} \cdot {I_0}^{\alpha}$, and thus appearing broader than both $W(f)$ and $I(f)$ by a factor $1/\sqrt{\alpha}$ (because $W(f)^{\alpha} = exp(-(f - f_c)^2/(2\sigma^2/\alpha)$ is a Gaussian of width $\sigma/\sqrt{\alpha}$ ). By the same reasoning as in the linear case, the ratio between the II tuning width measured at $L_R$ = 3 dB and the IR tuning width measured at $L_I$ (10 or 40 dB) would therefore be $2\sigma\sqrt{2ln(10^{3/10})}/\sqrt{\alpha}/2\sigma\sqrt{2ln(10^{L_I/10})} \approx \sqrt{\ln(2)/(\alpha \cdot \ln(10^{L_I/10}))}$. If instead, we assume that the compressive IO function precedes (rather than follows) frequency filtering, i.e., $R(f,I) = W(f) \cdot I^{\alpha}$, an analogous reasoning can be applied and the same ratio is obtained. In this case, it is the II tuning curve that is unaffected by the non-linearity (since $R(f) = W(f) \cdot {I_0}^{\alpha}$) and reflects the true transfer function $W(f)$, whilst the IR tuning curve is expanded ($I(f) = \sqrt[\alpha]{R_0}/\sqrt[\alpha]{W(f)}$) and thus appears sharper than both $W(f)$ and $R(f)$ by a factor $\sqrt{\alpha}$. Therefore, by the same reasoning as previously, the ratio between II tuning width at

3 dB and IR tuning width at $L_I$ (10 or 40 dB) is still $\sqrt{\ln(2)/(\alpha \cdot \ln(10^{L_I/10}))}$. Note that, because we do not know whether the non-linearity precedes or follows frequency filtering at the level of auditory cortical neurons, we do not know which of the IR or II tuning curves reflects the true transfer function, $W(f)$.

### Supplementary discussion

#### Relationship between fMRI adaptation tuning and population-average single-neuron response tuning

The neural response to a probe stimulus is suppressed when the probe is preceded by an adaptor stimulus. fMRI adaptation measures the average amount of suppression across the population of neurons contained within a given voxel or ROI. As a result of neuronal selectivity, the suppression (fMRI adaptation) will be tuned to the probe stimulus, as neurons more responsive to the probe will be more effectively stimulated – and thus adapted – by adaptors close to, rather than remote from, the probe. This does not mean, however, that population-average suppression selectivity is equal to population-average excitatory selectivity, because the population includes neurons that are not exactly tuned to the probe frequency, but nevertheless respond significantly to it. For these neurons, suppression is larger near their characteristic frequency than near the probe frequency, resulting in a widening of the population-average suppression tuning. Here we formally show that for a neuronal population with a wide range of characteristic frequencies, the widening corresponds exactly to a factor $\sqrt{2}$.

Under the assumptions that adaptation is multiplicative and proportional to the size of the adaptor response, the adapted probe response of a single neuron is given by $R_{cell}(P|A) = R_{cell}(P) \cdot (1 - \kappa \cdot R_{cell}(A))$, and thus the single-neuron adaptation amount, $R_{cell}(P) - R_{cell}(P|A)$, is given by the product $\kappa \cdot R_{cell}(P) \cdot R_{cell}(A)$. Here, $R_{cell}(P)$ is the unadapted probe response, $R_{cell}(A)$ is the response to the adaptor, and $\kappa$ is the neuronal adaptation strength (see main Methods section).

If the single-neuron responses, $R_{cell}$, are Gaussian-shaped, with spread $\sigma$, then the population-average adaptation amount across an unbiased neuron population (with neural characteristic frequencies spanning the entire frequency range, which, for simplicity, will be approximated by [0 ∞]) is given by

$$\int_0^\infty \kappa \cdot e^{-((f_P-x)^2+(f_A-x)^2)/(2\,\sigma^2)} dx = \sqrt{\pi}\,\kappa\,\sigma/2 \cdot e^{-(f_P-f_A)^2/(4\,\sigma^2)}\,\mathrm{erf}\left(\frac{2x-f_P-f_A}{2\,\sigma}\right)\Big|_0^\infty,$$

where $f_P$ and $f_A$ are the probe and adaptor frequencies, respectively and erf is the Gauss error function. Given that $\mathrm{erf}(\infty) = 1$, this simplifies to $\sqrt{\pi}\,\kappa\,\sigma/2 \cdot \mathrm{erf}\big((f_p + f_A)/(2\sigma)\big) \cdot e^{-(f_P-f_A)^2/(4\,\sigma^2)}$. Thus, the population-average adaptation amount is proportional to a Gaussian function, $e^{-(f_P-f_A)^2/(4\,\sigma^2)}$, of the adaptor frequency, $f_A$, with a peak at the probe frequency, $f_P$, and spread $\sqrt{2} \cdot \sigma$. This means that the widths of fMRI adaptation tuning curves would be expected to be wider by a factor of $\sqrt{2} \approx 1.4$, or 40%, than the underlying single-neuron response tuning curves.

### Supplementary tables and figures

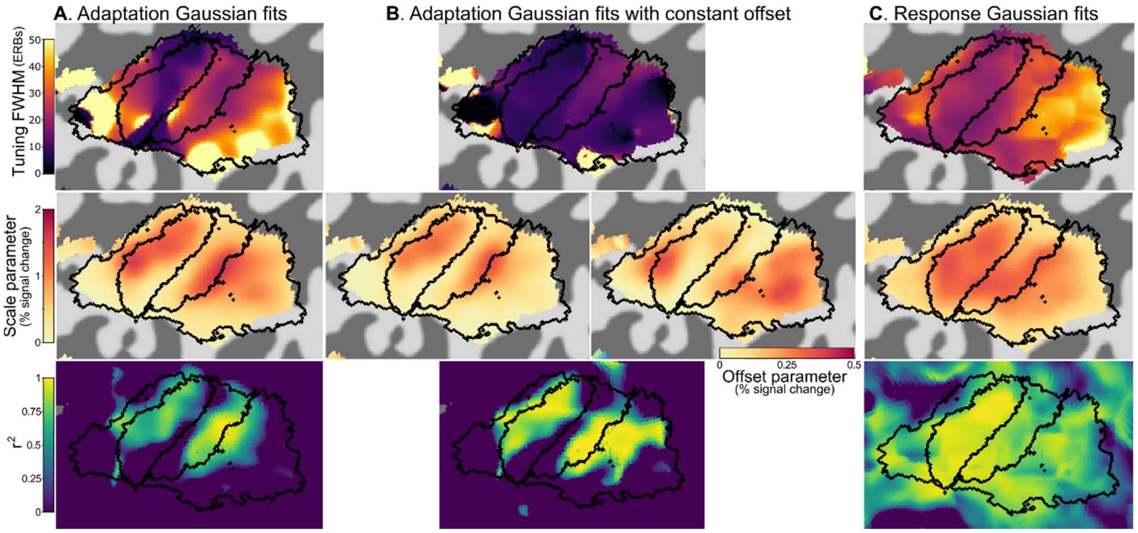


**Figure S1:** Gaussian parameter maps for searchlight fit of fMRI adaptation and BOLD response tuning widths (see methodological details in section A of the supplementary methods). A. FWHM, scale parameter and $r^2$ of simple Gaussian fit to voxelwise fMRI adaptation tuning curves. Black outlines represent anterior and posterior belt and core ROIs (see Fig. 2 in main document). Scale parameter and $r^2$ maps suggest that tuning curves of large enough amplitude to result in meaningful fits (large $r^2$) were largely restricted to the high-frequency-preferring regions at the borders between the anterior core and belt regions on the one hand and the posterior core and belt regions on the other hand. Nevertheless, FWHM maps show that, within these high-frequency regions, there were clear differences in tuning between the anterior core (most tuned), posterior core, anterior belt and posterior belt (least tuned) ROIs. B. FWHM, scale and offset parameter and $r^2$ of Gaussian fit with frequency-independent offset to voxelwise FMRI adaptation tuning curves. Regions with large $r^2$ values were similar to those for the simple Gaussian fits shown in A. Within these regions, tuning was generally narrower than in the simple Gaussian fit case, with less differences between different core and belt ROIs than in A. The offset parameter seemed larger in belt than core regions. Note that the generally larger $r^2$ values in B compared to A might be due to the Gaussian model with offset having an additional parameter. C. FWHM, scale parameter and $r^2$ of simple Gaussian fit to voxelwise BOLD response tuning curves. Compared to adaptation tuning curve fits shown in A and B, $r^2$ values were large in most parts of core and belt auditory cortex. Tuning was narrower in the anterior core ROI than in the posterior core ROI or anterior belt ROI, and was widest in the posterior belt ROI. Depending on the region, response tuning could be narrower or larger than the corresponding adaptation tuning shown in A.

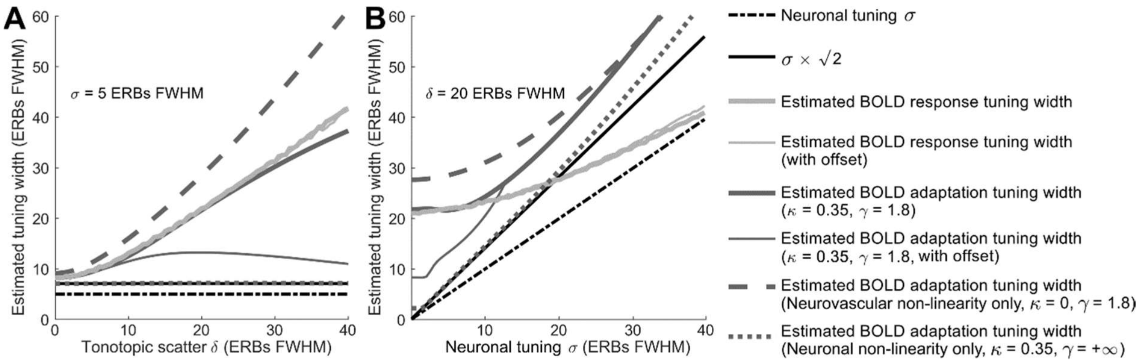


**Figure S2**: Estimated model BOLD response and fMRI adaptation tuning widths for a wide range (0 to 40 ERBs) of tonotopic scatter parameters (A) and neuronal tuning parameters (B). In both A and B, estimates from three versions of the model models are shown: a model only implementing neuronal adaptation ($\kappa$ = 0.35, $\gamma$ = +∞; dotted dark gray traces), a model only implementing neurovascular non-linearity ($\kappa$ = 0, $\gamma$ = 1.8; dashed dark gray traces) and a model implementing both ($\kappa$ = 0.35, $\gamma$ = 1.8; light and dark solid gray traces). Response tuning widths (light gray) and adaptation tuning widths (dark gray) were obtained by fitting either a simple Gaussian (thick gray traces) or a Gaussian with offset (thin gray traces). Response tuning width estimates are only shown for one type of model because they were identical for all three versions of the model (since these only depend on $\sigma$ and $\delta$). For models implementing either only neuronal adaptation or only neurovascular non-linearity, adaptation tuning estimates are only shown for one type of Gaussian fit because they were identical for both types of fit.

A. When neuronal tuning width is set to $\sigma$ = 5 ERBs FWHM (represented as the horizontal thin dashed-dotted black trace) and the tonotopic scatter parameter, $\delta$, varies from 0 to 40 ERBs, the response tuning width consistently overestimates the actual neuronal tuning width, and the amount of bias systematically increases with increasing tonotopic scatter, irrespective of whether it is estimated with a simple Gaussian (thick light-gray trace) or a Gaussian with offset (thin light-gray trace). Adaptation tuning width (solid dark gray traces) also systematically overestimates the neuronal tuning width, but the amount of bias depends on whether adaptation is caused by neuronal mechanisms or by neurovascular non-linearity: in the absence of neurovascular non-linearity, the adaptation tuning width obtained with a simple Gaussian (dotted dark-gray trace) overestimates neuronal tuning width by a constant factor of $\sqrt{2}$, irrespective of the amount of tonotopic scatter (compare with the thin solid back horizontal trace, representing $\sqrt{2} \times \sigma$ = 7.07 ERBs). Conversely, when adaptation is purely due to neurovascular non-linearity, the estimated adaptation tuning width (dark-gray dashed trace) strongly increases with increasing tonotopic scatter. For more realistic models using a combination of neuronal adaptation and neurovascular non-linearity, adaptation tuning width estimated using a simple Gaussian model (thick solid dark-gray trace) is intermediate between the previous two scenarios, and overestimation of the underlying neuronal tuning width increases overall with decreasing topographic organization. Attempting to measure the tuning width of only the frequency-specific adaptation component by fitting a Gaussian with frequency-independent offset (thin solid dark-gray trace) only partially reduces the bias, although this strategy becomes more successful for large values of the tonotopic scatter parameter (i.e. for non-topographically-organized cortical regions).

B. When the tonotopic scatter $\delta$ fixed at 20 ERBs and neuronal tuning width, $\sigma$, varies from 0 to 40 ERBs, similar biases are observed. Although the estimated response tuning width converges towards the true neuronal tuning width for larger values of $\sigma$ (i.e. the contribution of the tonotopic scatter becomes negligible for widely tuned neurons), adaptation tuning width overestimates the underlying neuronal tuning width at all tested values of $\sigma$. For purely neuronal adaptation, the discrepancy between actual neuronal tuning width and estimated adaptation tuning width again corresponds to a multiplicative constant of $\sqrt{2}$ at all values of $\sigma$ (or slightly greater for $\sigma$ > 20 ERBs). When introducing neurovascular non-linearity, the discrepancy increases substantially, particularly for small $\sigma$ values below about 20 ERBs. For these narrow neuronal tuning width values, fitting a Gaussian with frequency-independent offset reduces the bias, but does not eliminate it.

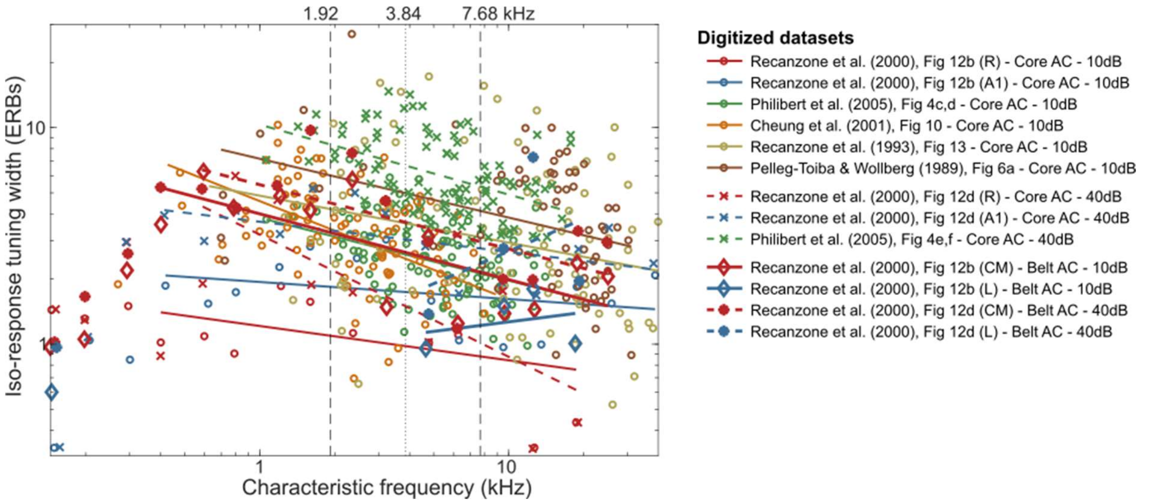


**Figure S3**: Non-human primate neuronal iso-response tuning width estimates as a function of neurons' characteristic frequencies, digitized from 13 datasets across five single- and multi-unit electrophysiological studies. Each point corresponds either to a single-/multi-unit or to the average of several single-/multi-units binned by characteristic frequency, depending on the study. Recordings were made either in core or belt auditory areas (small or large symbols), at intensity levels 10 dB or 40 dB (open circles/diamonds or crosses/filled circles). Tuning widths, originally given either in octaves or as quality factors, were converted to humans cochlear ERBs (see section B.1 in the supplementary methods section). Straight lines are best-fitting linear functions in log-log space for each dataset. In all except two datasets (both from belt area L), isoresponse tuning widths decreased with increasing characteristic frequency (at least for characteristic frequencies above 0.5 kHz). Vertical dashed lines indicate the range of characteristic frequencies we used to average tuning widths within datasets. For this figure, datasets were randomly decimated to a maximum of 100 points per dataset to avoid clutter.

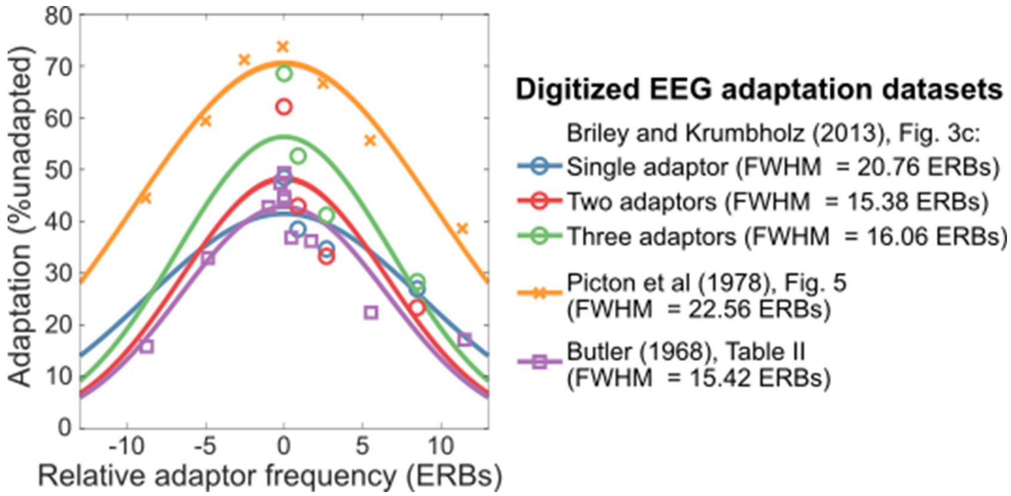


**Figure S4**: Frequency adaptation tuning curves from EEG studies. Each data point represents the group-average adaptation effect, computed as the decrease in N1 or N1-P2 ERP amplitude in response to a probe preceded by an adaptor, relative to the unadapted probe amplitude, expressed as a proportion. Adaptation is plotted as a function of the frequency difference between adaptors and probe. ERP adaptation tuning curves are plotted with best-fitting Gaussian functions whose FWHM is indicated in the legend.

**ROI-average fMRI adaptation tuning curves - Single-onset adaptor:**

| | Simple Gaussian model | | | | | | | Frequency-independent adaptation model | |
|---|---|---|---|---|---|---|---|---|---|
| | Outliers | FWHM (ERBs) | FWHM (oct) | Scaling | BIC | AICc | $r^2$ | BIC | AICc |
| Anterior Belt | 0 | 2180 ± 1819 | 179 ± 150 | 0.51 ± 0.32 | -68.9 | 69.4 | 0.00 | -72.8 | 67.3 |
| Anterior gradient | 3 | 22.1 ± 14.4 | 3.65 ± 2.36 | 0.75 ± 0.45 | -54.6 | 83.8 | 0.02 | -57.3 | 82.7 |
| Posterior Gradient | 3 | 22.3 ± 12.4 | 3.68 ± 2.04 | 0.65 ± 0.47 | -52.7 | 85.6 | 0.02 | -55.8 | 84.2 |
| Posterior Belt | 2 | 57.0 ± 28.7 | 9.58 ± 4.74 | 0.69 ± 0.29 | -104.1 | 34.2 | 0.02 | -107.0 | 33.0 |

**ROI-average fMRI adaptation tuning curves - Multiple-onset adaptor:**

| | Simple Gaussian model | | | | | | | Frequency-independent adaptation model | |
|---|---|---|---|---|---|---|---|---|---|
| | Outliers | FWHM (ERBs) | FWHM (oct) | Scaling | BIC | AICc | $r^2$ | BIC | AICc |
| Anterior Belt | 0 | 18.0 ± 7.7 | 2.97 ± 1.26 | 0.82 ± 0.24 | -109.2 | 57.1 | 0.27 | -94.3 | 74.0 |
| Anterior gradient | 0 | 9.0 ± 4.8 | 1.49 ± 0.79 | 0.74 ± 0.30 | -123.8 | 42.5 | 0.44 | -93.3 | 74.9 |
| Posterior Gradient | 0 | 12.9 ± 5.9 | 2.12 ± 0.97 | 0.82 ± 0.29 | -115.6 | 50.7 | 0.37 | -91.8 | 76.4 |
| Posterior Belt | 0 | 25.9 ± 5.6 | 4.28 ± 0.93 | 0.77 ± 0.22 | -144.9 | 21.4 | 0.33 | -124.7 | 43.5 |

**Gaussian model with asymptotic adaption offset**

| | Outliers | FWHM (ERBs) | FWHM (oct) | Scaling | Offset | BIC | AICc | $r^2$ |
|---|---|---|---|---|---|---|---|---|
| Anterior Belt | 0 | 10.0 ± 3.3 | 1.64 ± 0.55 | 0.69 ± 0.18 | 0.20 ± 0.17 | -108.0 | 56.4 | 0.31 |
| Anterior gradient | 0 | 11.0 ± 3.3 | 1.81 ± 0.54 | 0.82 ± 0.26 | -0.10 ± 0.15 | -120.1 | 44.3 | 0.44 |
| Posterior Gradient | 0 | 12.6 ± 4.9 | 2.07 ± 0.80 | 0.79 ± 0.30 | 0.03 ± 0.12 | -111.6 | 52.8 | 0.37 |
| Posterior Belt | 0 | 12.2 ± 3.2 | 2.01 ± 0.53 | 0.57 ± 0.16 | 0.26 ± 0.10 | -142.9 | 21.5 | 0.36 |

**ROI-average BOLD response tuning curves - Multiple-onset adaptor:**

| | Simple Gaussian model | | | | | | | Frequency-independent adaptation model | |
|---|---|---|---|---|---|---|---|---|---|
| | Outliers | FWHM (ERBs) | FWHM (oct) | Scaling | BIC | AICc | $r^2$ | BIC | AICc |
| Anterior Belt | 0 | 18.9 ± 3.2 | 3.40 ± 0.58 | 1.82 ± 0.36 | -391.5 | 180.9 | 0.75 | -94.3 | 74.0 |
| Anterior gradient | 0 | 16.7 ± 2.0 | 3.00 ± 0.35 | 1.71 ± 0.35 | -518.6 | 53.7 | 0.83 | -93.3 | 74.9 |
| Posterior Gradient | 1 | 19.8 ± 4.8 | 3.56 ± 0.85 | 1.57 ± 0.25 | -465.9 | 106.5 | 0.72 | -91.8 | 76.4 |
| Posterior Belt | 1 | 25.0 ± 8.1 | 4.53 ± 1.45 | 1.75 ± 0.19 | -268.2 | 304.1 | 0.55 | -124.7 | 43.5 |

**Gaussian model with asymptotic adaption offset**

| | Outliers | FWHM (ERBs) | FWHM (oct) | Scaling | Offset | BIC | AICc | $r^2$ |
|---|---|---|---|---|---|---|---|---|
| Anterior Belt | 0 | 18.6 ± 2.3 | 3.35 ± 0.41 | 1.88 ± 0.41 | -0.06 ± 0.16 | -387.2 | 181.9 | 0.75 |
| Anterior gradient | 0 | 17.2 ± 1.4 | 3.10 ± 0.25 | 1.75 ± 0.37 | -0.06 ± 0.06 | -517.7 | 51.4 | 0.83 |
| Posterior Gradient | 0 | 18.4 ± 2.6 | 3.31 ± 0.47 | 1.49 ± 0.30 | 0.08 ± 0.12 | -467.6 | 101.5 | 0.73 |
| Posterior Belt | 0 | 21.0 ± 3.6 | 3.79 ± 0.65 | 1.67 ± 0.34 | 0.15 ± 0.29 | -270.2 | 298.9 | 0.57 |

**Table S1:** Average parameter estimates and goodness-of-fit measures for models fitted to the individual-hemisphere ROI-average fMRI adaptation and/or BOLD response tuning curves. For participants in the single-onset adaptor condition (tables in the top row), fMRI adaptation tuning curves were fitted with either a simple Gaussian model (without frequency-independent adaptation offset parameter) or by a constant (frequency-independent) adaptation model (i.e. the mean adaptation across adaptor frequencies). For participants in the multiple-onset adaptor condition (tables in the 4 bottom rows), both fMRI adaptation and BOLD response tuning were fitted using three models: simple Gaussian, frequency-independent adaptation and a Gaussian model with asymptotic frequency-independent adaptation (offset parameter). For Gaussian models, average parameter estimates are given with their parametric 95% confidence intervals. The fitted Gaussian widths are given as half-maximum widths (FWHM) in both ERBs and transformed to approximate octave values (w.r. to the center of the adaptor frequency range for response tuning curves and to the probe frequency for adaptation tuning curves). The scale parameter refers to the best-fitting peak response in percent signal change for the response tuning curves, or the maximum normalized adaptation amount in percent for the adaptation tuning curves. For the second of the Gaussian fits, the offset parameter gives the best-fitting amount of frequency-independent adaptation/response, in the same units as the scaling parameter. The leftmost column in each table shows the number of rejected outliers for computing group-average parameters and for the ANOVAs testing for differences between ROIs. For the frequency-independent adaption model, the only estimated parameter was the average adaption value across all adaptor frequencies. Bayesian Information criterion (BIC), Akaike information criterion corrected for small sample size (AICc), and coefficient of determination ($r^2$) are for the group-fitted Gaussian models with respect to all individual-hemisphere tuning curves.

**Model comparison - Gaussian (tuned) vs frequency-independent adaptation**

| Single-onset adaptor (simple Gaussian) | | Multiple-onset adaptor: Simple Gaussian | | Multiple-onset adaptor: Gaussian with offset | |
|---|---|---|---|---|---|
| ΔBIC | ΔAICc | ΔBIC | ΔAICc | ΔBIC | ΔAICc |
| 3.9 | 2.2 | -35.2 | -10.8 | -21.5 | 6.7 |
| 2.8 | 1.1 | -62.8 | -38.4 | -36.0 | -7.8 |
| 3.1 | 1.4 | -55.8 | -31.4 | -36.0 | -7.8 |
| 2.9 | 1.2 | -35.9 | -11.6 | -17.7 | 10.5 |

**Simple Gaussian vs Gaussian with offset**

| Adaptation | | Response | |
|---|---|---|---|
| ΔBIC | ΔAICc | ΔBIC | ΔAICc |
| 1.2 | -0.7 | 4.3 | 1.1 |
| 3.7 | 1.8 | 1.0 | -2.3 |
| 3.9 | 2.1 | -1.7 | -5.0 |
| 1.9 | 0.1 | -1.9 | -5.2 |

**Table S2:** Model comparison between tuned (Gaussian) and frequency-independent adaptation (top tables), and between simple Gaussian and Gaussian with frequency-independent adaptation offset (bottom tables). ΔBIC and ΔAICc are computed by subtracting the BIC or AICc values for the more complex model (more parameters) from the values of the simpler model (with less parameters). Therefore, positive values for ΔBIC and ΔAICc indicate evidence for the simpler model and negative values indicate evidence for the more complex model. Absolute values of ΔBIC or ΔAICc can be interpreted as follows (Kass and Raftery 1995): 0 to 2 = weak evidence, 2 to 6 = positive evidence, 6 to 10 = strong evidence, >10 = very strong evidence.

| Study | Figure | Species | Anesthesia/task | Recording type | Region | Level | IR width | | II FWHM (α = 0.3) | | II FWHM (α = 0.1) | |
|---|---|---|---|---|---|---|---|---|---|---|---|---|
| | | | | | | | ERBs | Oct. | ERBs | Oct. | ERBs | Oct. |
| Recanzone et al. (2000) | Fig. 12 | Rhesus Monkey | Awake (behaving) | Single-unit | R | 10 dB | 1.1 | 0.18 | 1.1 | 0.18 | 1.9 | 0.31 |
| | | | | | | 40 dB | 2.0 | 0.33 | 1.0 | 0.16 | 1.7 | 0.28 |
| | | | | | A1 | 10 dB | 2.0 | 0.34 | 2.0 | 0.34 | 3.5 | 0.59 |
| | | | | | | 40 dB | 3.4 | 0.56 | 1.7 | 0.28 | 2.9 | 0.49 |
| Philibert et al. (2005) | Fig. 4 | Marmoset | Sodium pentobarbital | Multi-unit | A1 | 10 dB | 2.8 | 0.46 | 2.8 | 0.46 | 4.9 | 0.80 |
| | | | | | | 40 dB | 7.3 | 1.20 | 3.6 | 0.60 | 6.3 | 1.04 |
| Cheung et al. (2001) | Fig. 10 | Squirrel Monkey | Sodium pentobarbital | Both | A1 | 10 dB | 2.9 | 0.49 | 2.9 | 0.49 | 5.1 | 0.85 |
| Recanzone et al. (1993) | Fig. 13 | Owl Monkey | Sodium pentobarbital | Multi-unit | A1 | 10 dB | 6.4 | 1.05 | 6.4 | 1.05 | 11.1 | 1.82 |
| Pelleg-Toiba & Wollberg (1989) | Fig. 6 | Squirrel Monkey | Awake (passive) | Single-unit | A1 | 10 dB | 11.7 | 1.92 | 11.7 | 1.92 | 20.3 | 3.33 |
| Bartlett et la. (2011) | Fig. 4d | Marmoset | Awake (passive) | Single-unit | A1 | 10 dB | - | - | 1.5 | 0.26 | 1.5 | 0.26 |
| | | | | | | Best level | - | - | 2.3 | 0.38 | 2.3 | 0.38 |
| Kajikawa et al. (2005) | Fig. 7 | Marmoset | Ketamine hydrochloride | Multi-unit | A1 | 10 dB | 2.0 | 0.33 | 1.0 | 0.16 | 1.7 | 0.28 |
| | | | | | | 40 dB | 24.7 | 3.98 | 12.4 | 2.00 | 21.5 | 3.46 |
| | | | | | **Core (A1/R)** | | | ***Mean*** | ***5.0*** | ***0.8*** | ***8.5*** | ***1.4*** |
| | | | | | | | | *Median* | *2.8* | *0.5* | *4.9* | *0.8* |
| Recanzone et al. (2000) | Fig. 12 | Rhesus Monkey | Awake (behaving) | Single-unit | L | 10 dB | 1.0 | 0.16 | 1.0 | 0.16 | 1.7 | 0.27 |
| | | | | | | 40 dB | 1.4 | 0.22 | 0.7 | 0.11 | 1.2 | 0.19 |
| | | | | | CM | 10 dB | 2.9 | 0.48 | 2.9 | 0.49 | 5.0 | 0.84 |
| | | | | | | 40 dB | 4.1 | 0.68 | 2.1 | 0.34 | 3.6 | 0.59 |
| Kajikawa et al. (2005) | Fig. 7 | Marmoset | Ketamine hydrochloride | Multi-unit | CM | 10 dB | 18.7 | 3.02 | 18.8 | 3.03 | 32.5 | 5.24 |
| | | | | | | 40 dB | 30.0 | 4.82 | 15.0 | 2.41 | 26.1 | 4.18 |
| | | | | | **Belt (CM)** | | | ***Mean**** | ***6.3*** | ***1.03*** | ***10.8*** | ***1.76*** |
| | | | | | | | | *Median** | *3.5* | *3.48* | *6.0* | *6.03* |

**Table S3**: Average non-human primate tuning widths for each of 19 datasets recorded from different core and belt areas. Averages were computed either from the original iso-response tuning widths ("IR width" column), at the intensity level indicated by the "Level" column, or from tuning widths that were first converted to iso-intensity tuning curve FWHM ("II FWHM" columns) under different assumptions about the shape of the neuronal (and BOLD) input/output function, a power-law with a compressive exponent $\alpha$ equal either to 0.1 or 0.3 (see supplementary methods section B3). Corresponding mean and median values for core or belt auditory fields are reported in bold and italic respectively. *Belt mean and median values were corrected to account for the small number of belt datasets (see supplementary methods section B for details).

#### Supplementary references


Bartlett EL, Sadagopan S, Wang X. 2011. Fine frequency tuning in monkey auditory cortex and thalamus. J Neurophysiol 106:849-859.

Buus S, Florentine Meditors. Year Published|. Title|, Conference Name|; Year of Conference Date|; Conference Location| Place Published|:Publisher|. Pages p|.

Cheung SW, Bedenbaugh PH, Nagarajan SS, Schreiner CE. 2001. Functional organization of squirrel monkey primary auditory cortex: responses to pure tones. J Neurophysiol 85:1732-1749.

Eustaquio-Martin A, Lopez-Poveda EA. 2011. Isoresponse versus isoinput estimates of cochlear filter tuning. J Assoc Res Otolaryngol 12:281-299.

Glasberg BR, Moore BC. 1990. Derivation of auditory filter shapes from notched-noise data. Hear Res 47:103-138.

Hellman R, Zwislocki J. 1961. Some Factors Affecting the Estimation of Loudness. J Acoust Soc Am 33:687.

Kajikawa Y, de La Mothe L, Blumell S, Hackett TA. 2005. A comparison of neuron response properties in areas A1 and CM of the marmoset monkey auditory cortex: tones and broadband noise. J Neurophysiol 93:22-34.

Kass RE, Raftery AE. 1995. Bayes Factors. Journal of the American Statistical Association 90:773-795.

Lopez-Poveda EA, Eustaquio-Martin A. 2013. On the controversy about the sharpness of human cochlear tuning. J Assoc Res Otolaryngol 14:673-686.

Pelleg-Toiba R, Wollberg Z. 1989. Tuning properties of auditory cortex cells in the awake squirrel monkey. Exp Brain Res 74:353-364.

Philibert B, Beitel RE, Nagarajan SS, Bonham BH, Schreiner CE, Cheung SW. 2005. Functional organization and hemispheric comparison of primary auditory cortex in the common marmoset (Callithrix jacchus). J Comp Neurol 487:391-406.

Recanzone GH, Guard DC, Phan ML. 2000. Frequency and intensity response properties of single neurons in the auditory cortex of the behaving macaque monkey. J Neurophysiol 83:2315-2331.

Recanzone GH, Schreiner CE, Merzenich MM. 1993. Plasticity in the frequency representation of primary auditory cortex following discrimination training in adult owl monkeys. J Neurosci 13:87-103.

Uppenkamp S, Rohl M. 2014. Human auditory neuroimaging of intensity and loudness. Hear Res 307:65-73.

Wan X, Riera J, Iwata K, Takahashi M, Wakabayashi T, Kawashima R. 2006. The neural basis of the hemodynamic response nonlinearity in human primary visual cortex: Implications for neurovascular coupling mechanism. Neuroimage 32:616-625.
